# Engineered chimeras unveil swappable modular features of fatty acid and polyketide synthase acyl carrier proteins

**DOI:** 10.1101/2021.12.30.471467

**Authors:** Yae In Cho, Claire L. Armstrong, Ariana Sulpizio, Kofi K. Acheampong, Kameron N. Banks, Oishi Bardhan, Sydney J. Churchill, Annie E. Connolly-Sporing, Callie E.W. Crawford, Peter L. Cruz Parrilla, Sarah M. Curtis, Lauren M. De La Ossa, Samuel C. Epstein, Clara J. Farrehi, Grayson S. Hamrick, William J. Hillegas, Austin Kang, Olivia C. Laxton, Joie Ling, Sara M. Matsumura, Victoria M. Merino, Shahla H. Mukhtar, Neel J. Shah, Casey H. Londergan, Clyde A. Daly, Bashkim Kokona, Louise K. Charkoudian

## Abstract

The strategic redesign of microbial biosynthetic pathways is a compelling route to access molecules of diverse structure and function in a potentially environmentally sustainable fashion. The promise of this approach hinges on an improved understanding of acyl carrier proteins (ACPs), which serve as central hubs in biosynthetic pathways. These small, flexible proteins mediate the transport of molecular building blocks and intermediates to enzymatic partners that extend and tailor the growing natural products. Past combinatorial biosynthesis efforts have failed due to incompatible ACP-enzyme pairings. Herein we report the design of chimeric ACPs with features of the actinorhodin polyketide synthase ACP (ACT) and of the *E. coli* fatty acid synthase (FAS) ACP (AcpP). We evaluate the ability of the chimeric ACPs to interact with the *E. coli* FAS ketosynthase FabF, which represents an interaction essential to building the carbon backbone of the synthase molecular output. Given that AcpP interacts with FabF but ACT does not, we sought to exchange modular features of ACT with AcpP to confer functionality with FabF. The interactions of chimeric ACPs with FabF were interrogated using sedimentation velocity experiments, surface plasmon resonance analyses, mechanism-based crosslinking assays, and molecular dynamics simulations. Results suggest that the residues guiding AcpP-FabF compatibility and ACT-FabF incompatibility may reside in the loop I, α-helix II region. These findings can inform the development of strategic secondary element swaps that expand the enzyme compatibility of ACPs across systems and therefore represent a critical step towards the strategic engineering of ‘unnatural’ natural products.

## Introduction

Natural product biosynthetic pathways generate an astounding array of structurally diverse, and often complex, organic molecules. In nature, these molecules can serve roles essential to the survival of an organism (‘primary metabolites,’ such as fatty acids) or can mediate inter-organismal relationships through the conveyance of competitive advantages (‘secondary metabolites’, such as antibiotics). From a public health perspective, natural products have played a pivotal role in drug discovery for the treatment of cancer, infection, and cardiovascular disease.^1^ While structural complexity creates barriers to obtaining natural products and their analogs via total synthetic approaches, recent technological and scientific advances pave the way to gain access to expanded chemical diversity via genome mining and biosynthetic engineering.^2–4^ Understanding how microorganisms manufacture natural products on a molecular level is a critical step towards more fully harnessing this remarkable biosynthetic machinery to benefit human health and the environment.

Fatty acids and polyketides are two classes of natural products that are manufactured by teams of enzymes encoded by biosynthetic gene clusters.^5,6^ The enzyme assemblies (termed ‘synthases’) are classified as type I or type II depending on whether the synthase is comprised of polypeptides with multiple domains or numerous stand-alone proteins, respectively. In type II fatty acid synthases (FASs) and polyketide synthases (PKSs), a single acyl carrier protein (ACP) is responsible for presenting malonyl-based building blocks and biosynthetic intermediates to the other members of the synthase. These biomolecular interactions proceed with remarkable fidelity and efficiency (**Figure 1**).

**Figure 1.**
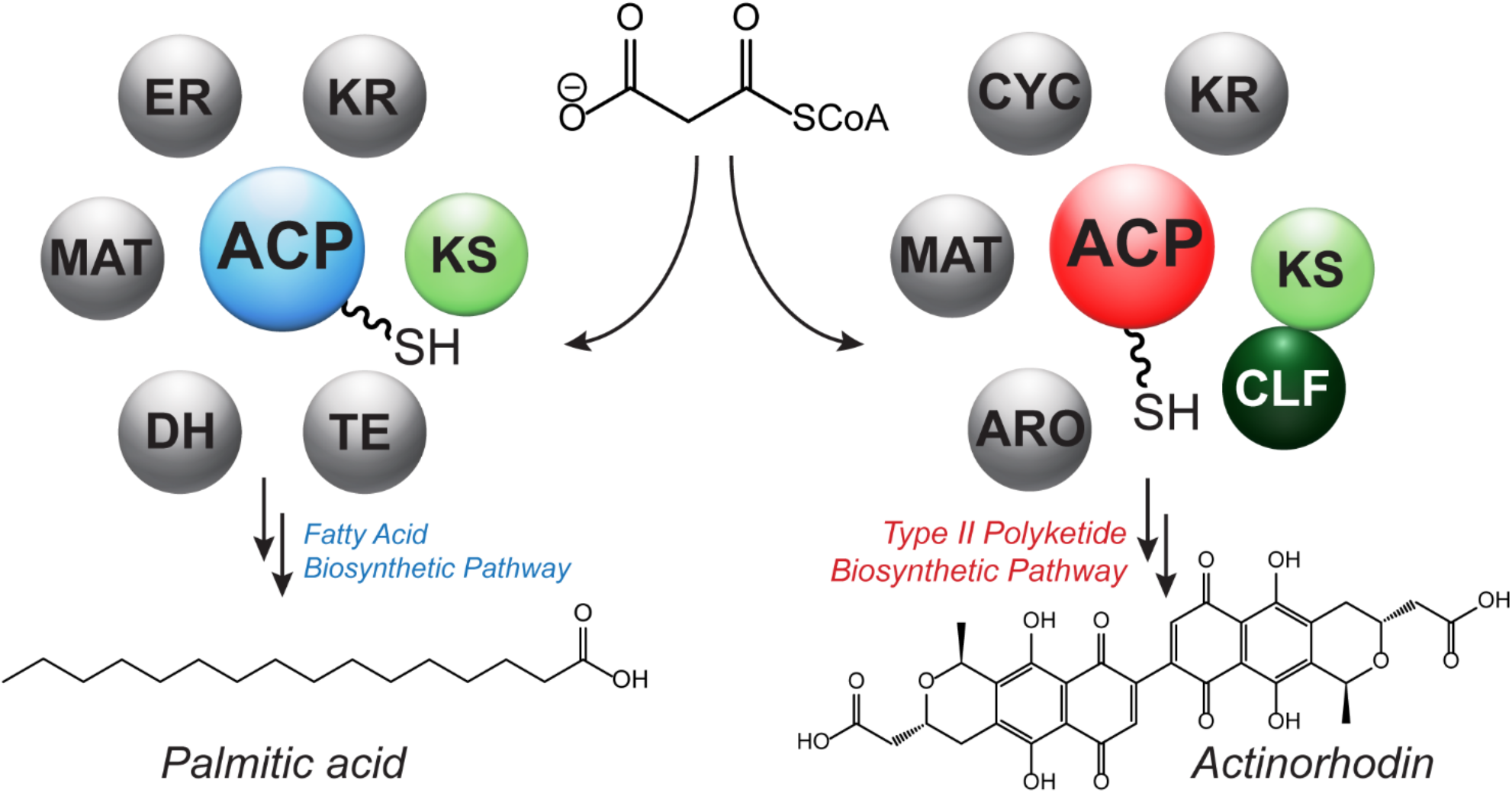
ACPs serve as a central hub in type II polyketide and fatty acid biosynthetic pathways. In type II fatty acid synthases (FASs) the ACP interacts with a suite of enzymes to manufacture fatty acid products (left). In type II polyketide synthases (PKS) the ACP interacts with suite of enzymes to manufacture polyaromatic polyketide products (right). Despite their similar structures and functions, type II FAS and type II PKS ACPs are not interchangeable. Abbreviations: ACP = acyl carrier protein, ARO = aromatase, CLF = chain length factor, CYC= cyclase, DH = dehydratase, ER = enoyl reductase, KR = ketoreductase, KS = ketosynthase, MAT = malonyl-CoA:ACP transacylase, TE = thioesterase.

These small, ∼8 kDa proteins are post-translationally modified into their active ‘*holo*’ forms which includes a flexible 18 Å 4’-phosphopantetheine (Ppant) arm appendage that acts to bind and carry molecular cargo between cognate enzymatic partners. The conformationally dynamic nature of the Ppant arm is thought to play a critical role in their versatility within a given synthase (**Figure 2**).^7–9^

**Figure 2.**
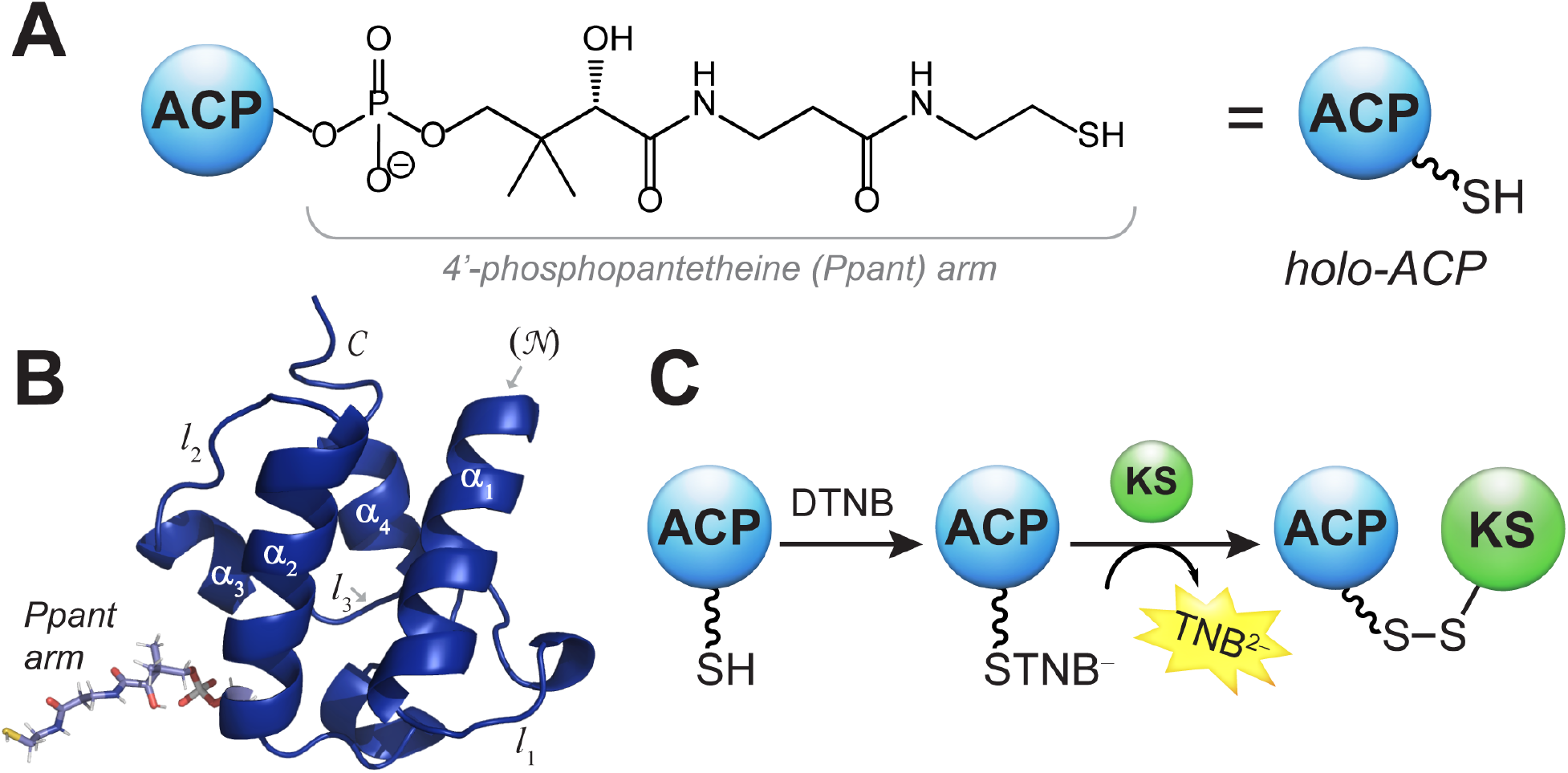
Type II PKS and FAS ACPs display conserved structural features, and their mechanistically relevant binding to the *E. coli* KS FabF can be evaluated via a colorimetric assay. (A) ACPs are converted to their active “*holo*” forms via the installation of a coenzyme A-derived phosphopantethiene (Ppant) arm. (B) The ACP (PDB: 2FAC) features four α-helices connected by unstructured loops. (C) In this study, the interactions of ACPs with the ketosynthase (KS) from the fatty acid biosynthetic pathway were monitored via the installation of a reactive colorimetric probe, as outlined previously.^10^

The overall structure and function of ACPs are largely conserved across type II FASs and PKSs.^7– 9,11–16^ These commonalities are supported by the inference that type II PKSs and FASs share a common ancestor, despite the distinct structures and biological functions of their molecular outputs (**Figure 1**).^17^ Interestingly, while some interchangeability has been observed within FAS ACPs *in vivo*,^18^ several *in vitro* mechanism-based crosslinking studies have revealed a lack of interchangeability between type II FAS and PKS ACPs. Take for example the ACP-KS interaction, which facilitates building the carbon backbone of the natural product through repeated chain extension (decarboxylative condensation) and chain translocation (transacylation) reactions in type II PKSs and FASs: Multiple studies have shown that despite its structural similarities to the *E. coli* FAS ACP (AcpP), the ACP from the actinorhodin (ACT) type II PKS does not interact with the FabF ketosynthase (KS) from the *E. coli* type II FAS.^10,19–22^ The lack of interchangeability of ACPs across type II FASs and PKSs creates a barrier to redesigning synthases to general novel natural products.

In theory, mixing-and-matching enzymes from different pathways could give rise to large libraries of novel compounds by converging the diversity programmed by essential enzymes across synthases (*e*.*g*. the range of carbon backbone chain length) with the varied oxidation states and cyclization patterns directed by accessory enzymes (*e*.*g*. ketoreductions, intramolecular aldol-condensations, etc.). However, this reality can only be realized if ACPs can be strategically engineered to interact with non-cognate enzymatic partners. While mutagenesis could enable individual pathway engineering, a modular approach in which entire secondary features are swapped lends itself more readily to streamlined combinatorial biosynthetic approaches.^23–25^ Therefore, uncovering the modular elements of the type II ACPs that direct their compatibility within non-cognate synthases represents a critical step towards effective combinatorial biosynthesis.^7,8^

As part of a course-based undergraduate research experience (CURE), we sought to uncover the modular elements that dictate the compatibility between ACPs and KSs, a partnership that is essential to a functioning synthase. More specifically, we investigated the molecular underpinnings of AcpP and FabF compatibility and ACT and FabF incompatibility. Taking a student-led divide-and-conquer approach, we designed 16 ACP chimeras (two per student pair), each of which harbored modular elements of AcpP and ACT (**Figure 3**). Following several studies that pointed towards helix II as playing an important role in ACP-protein recognition,^26–30^ we designed several chimeras aimed to parse out the role of this particular region. Students developed their biochemistry laboratory skills while designing chimeric ACP expression plasmids, assembling these plasmids via Gibson Assembly,^31^ and expressing, purifying, and characterizing the chimeric ACPs. Results from these experiments, coupled with molecular dynamics (MD) simulations, pointed to the linker 1 and helix II region as the modular element responsible for ACT incompatibility and AcpP compatibility with FabF. This work lays a foundation for building more robust hybrid combinatorial synthases capable of manufacturing novel chemical diversity.

**Figure 3.**
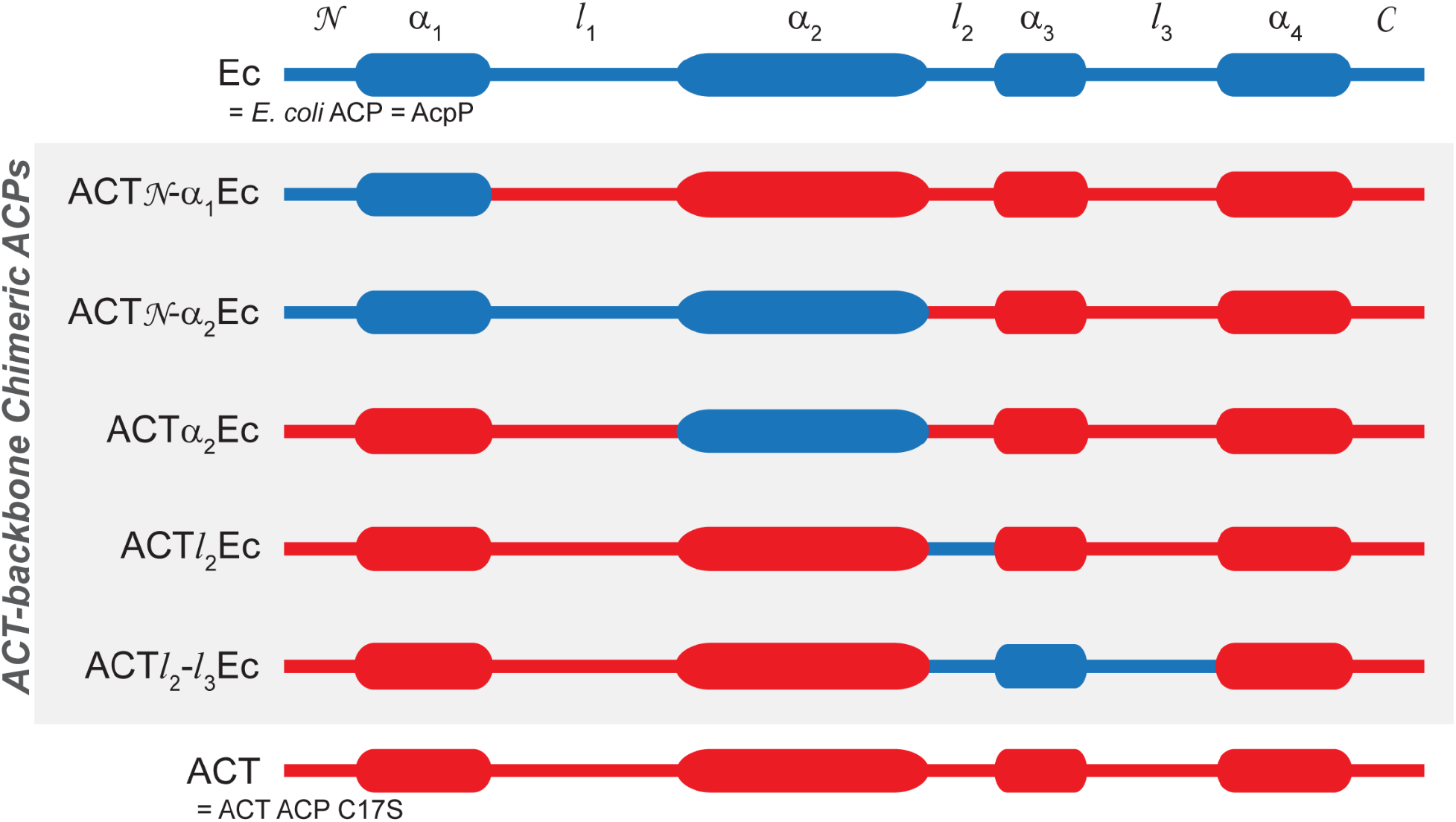
Acyl carrier protein (ACP) chimeras used in this study. The ACP chimeras are depicted with blue representing AcpP sequences and red representing ACT (C17S mutant) sequences. The chimeras shown in grey box are with ACT backbones and AcpP (“Ec”) insets. Not pictured here, but other chimeras were also studied with the same modular swaps, but with AcpP (“Ec”) backbones and ACT insets (**Table S1** and **S2**).

## MATERIALS AND METHODS

### Construction, expression, and purification of AcpP/ACT chimeras

Each pair of students designed and constructed two complementary ACP chimeras: one with an ACT-backbone and one AcpP-backbone. In total, the eight ACT-backbone and eight AcpP-backbone chimeras (**Figure 3**) were modeled using Phyre2^32^ (**Table S1**) and constructed using Gibson Assembly (**Table S2**) as described in Supporting Information. AcpP, ACT,^10^ and the chimeric ACPs were transformed into competent BAP1 *E. coli* cells which contained a phosphopantetheinyl transferase to convert the *apo* ACPs into their *holo* forms.^33^ pXY-FabF, kanamycin-resistant, was transformed into *E. coli* BL21 cells. ACPs were expressed and purified as described in the Supporting Information. Owing to *E. coli* harboring a native ACP phosphodiesterase, some of the expressed ACPs were isolated as *apo* and *holo* mixtures.^34^ An additional, promiscuous, phosphopantetheinyl transferase Sfp-catalyzed reaction was used to convert the remaining *apo* ACPs into 100% *holo*.^35^ Prior to applying the eluant onto a poly-prep (Bio Rad) column, 200 μL of expressed ACPs equilibrated resin was removed for analysis by LC-MS to evaluate the ratio of *apo*-vs *holo*-ACP in solution.^12^ When necessary, ACPs were converted to 100% *holo* using an on-resin Sfp reaction, as described in Supporting Information.^12^ FabF (KS) was purified as described previously.^10^ All proteins were analyzed by SDS PAGE as described in Supporting Information.

### Circular Dichroism (CD)

Circularly dichroism (CD) spectra were acquired to assess the secondary structure of chimeric *holo-* and TNB-ACPs. The helical compositions of the ACP chimeras were compared to that of wild type AcpP and ACT. Samples of 0.5 mg/mL ACPs were loaded into a 1 mm quartz cuvette (Hellman Analytics) and CD wavelength scans between 180–260 nm recorded on an Aviv model 410 spectropolarimeter. All samples were collected at 25 °C using a 1 nm bandwidth, a step resolution of 0.5 nm, and 3 seconds averaging time. The baseline was corrected against the storage buffer between each run. Data were normalized and plotted in Origin (Version 8.60, OriginLab Corporation, Northampton, MA, USA) (**Figure S14**). Changes in signal at 222 nm were followed as a function of temperature using the following parameters: ∼10–90 °C, 2 °C steps, 2-minute equilibrium, heating rate of 2 °C min^−1^, 30 second signal averaging time, and a 1 nm bandwidth. Data were analyzed via CDpal using a two-state unfolding model N⇌U with the standard assumption that ΔC_p_ = 0, as described previously.^12^

### Conversion of *holo-*ACP to ACP-TNB^−^

For each ACP, the conversion of the terminal thiol of the ACP Ppant arm into the mixed disulfide ACP-TNB^−^ was achieved upon reacting the *holo* ACP (0.5–1 mM in 50 mM sodium phosphate buffer, pH 7.6) with 8 molar equivalents of 25 mM DTNB in 50 mM sodium phosphate buffer, pH 7.6, for one hour at room temperature in a microfuge tube. The formation of the mixed disulfide ACP-TNB^−^ species was observed via a color change to bright yellow, indicating the release of TNB^2–^. The ACP-TNB^−^ was isolated and desalted using a Sephadex G-25 PD-10 desalting column (GE Healthcare, #17085101) equilibrated with 50 mM sodium phosphate buffer, pH 7.6. The proteins were flash frozen and stored in –80 °C in 200 μL aliquots at a concentration of ∼200–300 μM.

### Liquid Chromatography-Mass Spectrometry (LC-MS)

ACP samples (20 μL, ∼50 μM in 50 mM sodium phosphate, pH 7.6) were analyzed by LC-MS to determine the *apo/ holo* ratio and to confirm the successful attachment of TNB^2–^ to *holo* ACP. A sample volume of 10 μL was injected into a Waters XBridge Protein BEH C4 Column (300A, 3.5 μM, 2.1 mm x 50 mm) heated to 45 °C for analysis by electrospray ionization mass spectrometry (ESI MS) in the positive mode. Protein samples were eluted using the following conditions: 0–1 min 95% A, 3.1 min 5%, 4.52 min 5% A, and 4.92–9 min 95% A (Solvent A = 99.9% (v/v) water + 0.1% (v/v) formic acid; Solvent B = 99.9% (v/v) acetonitrile + 0.1% (v/v) formic acid). The acquired mass spectra were deconvoluted using ESIprot^36^ online and plotted using Origin (Version 8.60. OriginLab Corporation, Northampton, MA, USA). The observed and calculated molecular weights for *apo, holo*, and TNB^−^ attached ACPs were compared to verify the presence of the desired moiety (**Figure S1**–**S9**).

### Crosslinking Assays: ACP-TNB + FabF

A colorimetric crosslinking assay was used to determine whether or not chimeric ACPs formed a mechanism-based crosslink with FabF, using methods similar to those previously described.^10^ ACP-TNB^−^chimeras, AcpP-TNB^−^ and ACT-TNB^−^ were each incubated with FabF in 50 mM sodium phosphate buffer, pH 7.6, for 20 min at room temperature. A gradient of ACP-TNB^−^: FabF molar ratios (1:0, 0:1, 0.2:1, 0.5:1, 1:1, 1.5:1, and 2:1) were used to assess any molar-dependent changes in formation of a crosslinked species. The ACP concentration was altered relatively to the FabF concentration, with FabF held constant at 25 μM. The samples were analyzed by SDS PAGE, UV-Vis spectroscopy, and size exclusion chromatography (**Figure 4** and **Figures S15** and **S16**).

**Figure 4.**
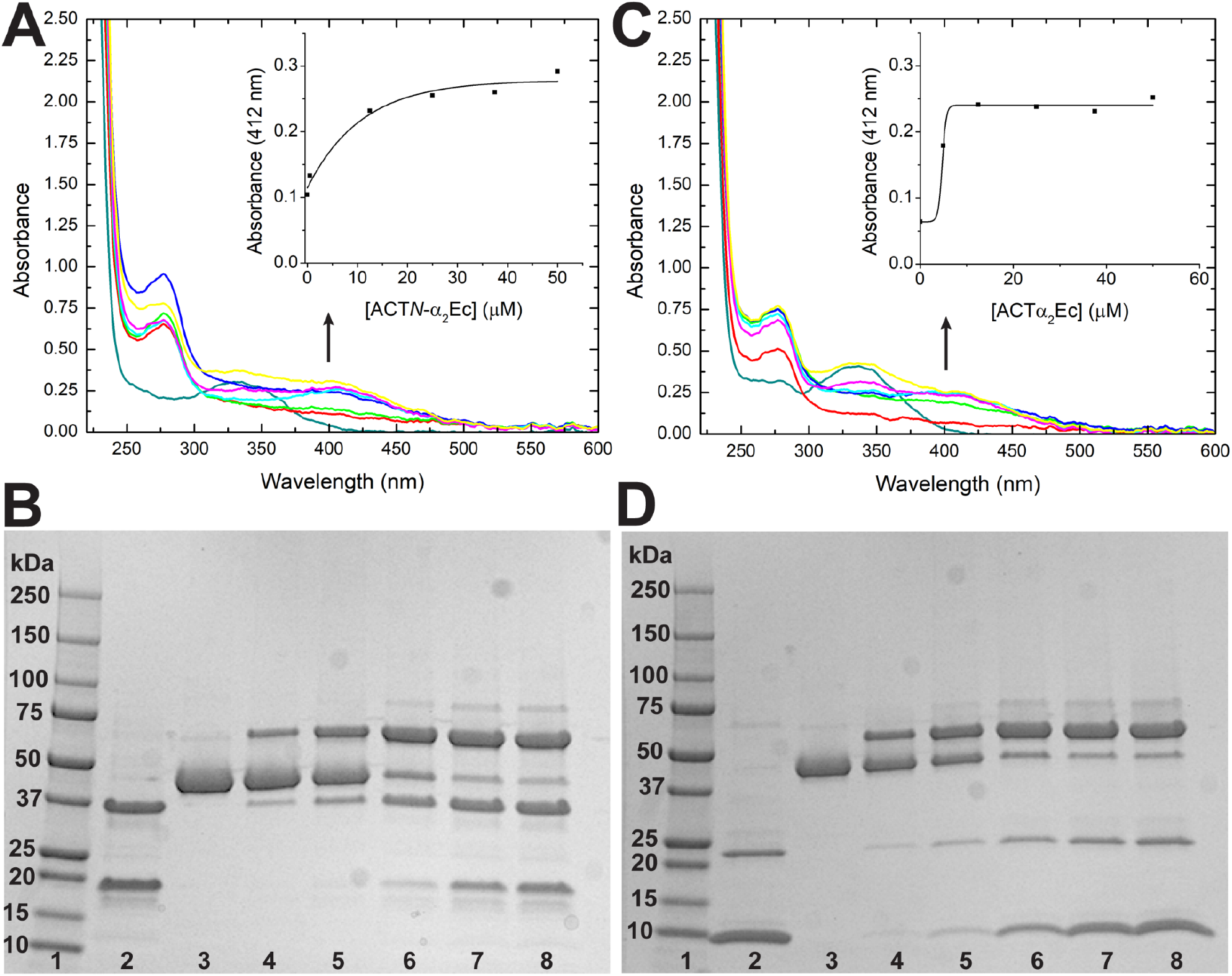
Mechanistic crosslinking studies indicate that ACT𝒩-α_2_Ec and ACTα_2_Ec form a complex with the *E. coli* KS FabF. Increase in chimera ACP concentrations corresponds with an increase in the concentration of the crosslinked complex. The concentration of FabF was kept at 25 μM, and ACT𝒩-α_2_Ec (A, B) and ACTα_2_Ec (C, D) varied from 0–50 μM. Legend for panels A and C: dark cyan for ACP chimera alone (5 μM); red for FabF (25 μM); green for ACP chimera (5 μM) + FabF (25 μM); blue for ACP chimera (12.5 μM) + FabF (25 μM); cyan for ACP chimera (25 μM) + FabF (25 μM); magenta for ACP chimera (37.5 μM) + FabF (25 μM); yellow for ACP chimera (50 μM) + FabF (25 μM). For panels B and D lanes 1–8 represent: 1. Protein ladder; 2. ACP chimera alone (5 μM); 3. FabF (25 μM); 4. ACP chimera (5 μM) + FabF (25 μM); 5. ACP chimera (12.5 μM) + FabF (25 μM); 6. ACP chimera (25 μM) + FabF (25 μM); 7. ACP chimera (37.5 μM) + FabF (25 μM); 8. ACP chimera (50 μM) + FabF (25 μM).

### Size Exclusion Chromatography (SEC)

All SEC samples were mixed with 20 mM Tris-HCl, 150 mM NaCl pH 7.0, to achieve a total volume of 500 μL. The samples were injected and run through a ÄKTA Pure chromatography gel filtration system (GE Healthcare Life Sciences) equipped with a Superdex 75 Increase 10/300 GL column (GE Healthcare Life Sciences) with a molecular weight range of 3000-7000 Da equilibrated in 20 mM Tris-HCl, 150 mM NaCl, pH 7.0. 1 mL fractions were eluted from 1.50 column volumes of the same buffer at a flow rate of 0.8 mL/min. Chromatograms were analyzed using the associated UNICORN™ software (Version 7.1, Cytiva, Marlborough, MA) package and plotted using Origin (Version 8.60. OriginLab Corporation, Northampton, MA, USA).

### Sedimentation Velocity with the Analytical Ultra Centrifuge (SV-AUC)

Sedimentation velocity experiments were performed using Beckman new model Optima Analytical Ultracentrifuge equipped with an An-50 Ti rotor. Samples were dialyzed overnight against 20 mM Tris-HCl, pH 8.0, 150 mM NaCl, and 1 mM DTT and loaded onto two-channel Epon, charcoal-filled centerpieces with 1.2 cm path length containing 415 μL sample and 430 μL dialysis buffer. Sedimentation boundaries were measured at a speed of 42,000 rpm until samples cleared the cell. All measurements were performed at 20 °C using a step size of 0.002 cm and time delay of 0 s. Samples were monitored at 250 nm and 280 nm with a required starting absorbance between 0.3 and 1.0. Data collected at 280 nm were used in data analysis.

The FabF concentration-dependence was established in our previous published data. For titration experiments, the concentration of FabF was kept constant at 17 μM and ACTα_2_Ec concentrations varied 0–50 μM. Sedimentation coefficient, c(s) distributions were generated using Sedfit (v.15.3)^37^ using continuous c(s) distribution model and only Time Independent (TI) Noise was fitted. The c(s), distributions were plotted utilizing Gussi (v.1.3.2)^38^ and the effective particle theory (EPT) fast isotherm was constructed by defining the reaction boundary to be between 4 and 8 S, and for the s(w) isotherm integration of the c(s) distribution between 1–8 S defining the exclusion zone for the isotherm between 2–4 S. EPT s(w) fast and s(w) isotherms were globally fit to an A + B ↔ AB hetero-association implemented in Sedphat (v.14.0).^39^ The sB, sAB, of the complex were fixed at respective 1.7 S, and 6.6 S while the sAB was fitted parameter.

The binding affinity of ACTα_2_Ec to FabF was determined by applying the weighted-average sedimentation coefficient (s_w_) as a function of ACTα_2_Ec concentration. Partial saturation of boundary was achieved by titrating 17 μM FabF with up to 50 μM ACTα_2_Ec. The effective particle theory (EPT) fast isotherm was constructed by defining the reaction boundary between 4 and 8 S, and a second isotherm was constructed by integrating the c(s) distribution between 1 and 8 S defining the exclusion zone between 2 and 4 S. The binding strength of 140 μM was obtained when both isotherms were fitted to an A + B ↔ AB model by varying sA and fixing both sB and sAB respectively at 1.7 S and 6.6 S. The estimated K_D_ of 140 μM indicates a much weaker binding for the ACTα_2_Ec to FabF when compared to that of the wild type AcpP.^10^

### Open Surface Plasmon Resonance (OpenSPR)

The binding affinities (K_D_) of the ACPs to FabF KS were analyzed using the OpenSPR instrument (2-Channel) and NTA sensor chips from Nicoya Lifesciences. All protein samples were SEC purified prior to the SPR experiments using the buffer (20 mM Tris-HCl, pH 7.0 with 150 mM NaCl that was filtered (0.22 μm) and degassed overnight. For each SPR experiment, a NTA sensor chip was freshly ligand-immobilized. The sensors were reused only up to two experiments. The instrument setting for experiment temperature was kept at 20 °C. All running buffers and reagents were either kept or brought up to r.t. before use. All protein samples were thawed, prepared, and kept on ice until the injection. For every injection, a minimum of 150 μL of sample (with no visible air bubble) was injected into both channels, unless mentioned otherwise. Two types of running buffers were used: PBST (phosphate buffered saline with 0.05% (w/v) TWEEN-20), and PBST + 1% (w/v) BSA (bovine serum albumin), both of which were filtered (0.22 μm) prior to use. The instrument was first primed with PBST, which was used as the running buffer for i) surface activation, ii) ligand immobilization, and iii) blocking. Then, the instrument was re-primed (buffer-exchanged) with PBST + 1% BSA before the analyte injections. The analytes (ACPs) were prepared by a serial dilution using PBST + 1% BSA (3-fold dilution from 150 μM to 2.54 nM for *holo*-AcpP and *holo*-ACTα_2_Ec; 3-fold dilution from 300 μM to 1.69 nM for *holo*-ACT).

For activating the NTA sensor surface with Ni^2+^, the injections were performed in the following order: 3 × 10 mM HCl (150 μL/min) → 2 × 350 disodium EDTA, pH 8 (100 μL/min) → 40 mM NiCl_2_ (20 μL/min). For ligand immobilization, 1 μM His-FabF KS (from a freshly thawed aliquot and diluted with PBST) was injected only to Channel 2 at 10 μL/min. Then, 10 μg/mL His-streptavidin (diluted with PBST) was injected at 10 μL/min through both channels to block any un-immobilized spots on the sensor chip.

For each analyte injection, flow rate was kept at 50 μL/min. Before the next analyte injection, a 10-min off-time was allowed for a complete dissociation. Binding affinity parameters were obtained by fitting the collected data to the “Affinity (a one-to-one binding model)” model using the TraceDrawer software (1.9.1).

### ACP-KS Complex Modeling

The amino acid sequences of ACT and AcpP were submitted to the I-TASSER server for structural modeling by protein fold recognition.^40^ The model given under the “Top 5 final models predicted by I-TASSER” heading with the highest C-score (typically Model 1) was chosen as the structural model for the ACP of interest. The structural model for FabF (PDB access code: 2GFW) was downloaded from the Protein Data Bank (PDB). After retrieval of the ACP and KS models of interest from I-TASSER, the .pdb files of the ACPs and the FabF homodimer were opened in PyMOL with an experimentally determined crosslinked crystal structure of FabF with AcpP (PDB access code: 6OLT) for alignment.^41^ This was repeated for FabF in contact with both ACPs. After alignment with the PDB structure 6OLT in PyMOL, the *apo*-ACPs (which result from I-TASSER’s sequence-only prediction) were converted to *holo*-ACPs by manual replacement with the “PNS” residue, an object that replicates the covalently attached Ppant arm at the serine attachment to each ACP.

### Molecular Dynamics Simulations

The ACP-KS-CLF structural models were prepared for larger-scale MD simulations on local hardware using GROMACS 2020.1. Equilibration and production runs were performed on Stampede2, a supercomputing cluster at the Texas Advanced Computing Center (TACC), in GROMACS 2018.3 on eight Knights Landing (knl) nodes with 64 Message Passing Interface (MPI) processes each (for a total of 512 MPI processes). All simulations used the CHARMM36 forcefield, with the artificial residue PNS (which includes the covalently linked Ppant arm) with parameters derived from CGenFF.^42^ The complex was centered in a cubic box with a distance of at least 1.0 nm from the complex to the edges of the box. The box was solvated with water (TIP3P) and the protein complex was neutralized with Na^+^ or Cl^-^ ions. After two 500 ps initial equilibration steps, 1 μs total time production simulations were run with 2 fs time steps.

General analysis of the trajectory files was performed using the Python MDAnalysis package. Solvent-accessible surface area (SASA) for the Ppant arm was evaluated using built-in functionality in Gromacs, and salt bridges were determined for each frame using MDAnalysis (see the SI file for more details). Statistical differences between the RMSD, SASA, and salt bridge data from different ACP-KS pairs were compared quantitatively using a two-tailed Student’s *t*-test.

### Free energy calculations

Free energy calculations were performed for five ACP structures: AcpP, ACT, Ecα_2_ACT, ACT 𝒩 - α_2_Ec, and ACTα_2_Ec, using the python-based toolkits OpenMM, mdtraj, and pymbar.^42–44^ Simulations leading to free energy calculations were initiated from the same docked structures described above and using the same MD force fields. Following equilibration, the systems were prepared for umbrella sampling. 10 systems were prepared where the center of mass distance between the ACP and KS was held still with a 0.1 kcal/mol/Å^2^ harmonic force. In each simulation, the complex was held to a different distance between 30 and 60 Å, evenly spaced between the 10 systems. After 2 ns of NVT (300 K) equilibration, the systems were run in NVT for 60 ns, collecting samples every 300 ps. The center of mass distance between the KS and the ACP was calculated for every sample, and the multistate Bennett acceptance ratio method was used to calculate the free energy differences between the bound states.^43^ These free energy differences were then used to calculate a potential of mean force along the center of mass distance coordinate.^45,46^ Please see the SI file for the full methods, results and discussion of these free energy calculations.

## RESULTS AND DISCUSSION

### Construction, expression, and purification of AcpP/ACT chimeras

A total of 16 plasmids encoding for the expression of *N*-terminally His_6_-tagged ACT/AcpP ACP chimeras were designed and constructed by the 16 students enrolled in a biochemistry CURE at Haverford College. Each student pair designed two chimeric ACPs: one with ACT backbone and the other with AcpP (“Ec”) backbone, each with the complementary inset at the same region on the sequence (**Table S1** and **S2**). We chose to express and characterize ACP chimeras that spanned the key structural elements of the ACP: alpha helices 1–4 (α_1_-α_4_), loops 1-3 (*l*_1_-*l*_3_) and unstructured *N*- and *C*-terminal regions. For target ACT-backbone-chimeras, elements of the *E. coli* FAS AcpP sequence were swapped into the corresponding region of the PKS actinorhodin ACT C17S mutant^10^ (*e*.*g*. ACTα_2_Ec represents the ACT C17S mutant^10^ with helix II replaced by the corresponding AcpP sequence). For target AcpP-backbone-chimeras, elements of the ACT (the C17S mutant) were swapped into the corresponding region of the AcpP (*e*.*g*. Ecα_2_ACT represents AcpP with helix II replaced by the corresponding ACT C17S sequence). All ACP chimeras were analyzed using the protein fold recognition server Phyre2^32^ and constructed via Gibson Assembly.^31^

Initial studies focused on the expression and purification of the eight ACT-backbone ACP chimeras. Despite expressing the chimeric ACP plasmids in BAP1 cells,^33^ which harbor the phosphopantetheinyl transferase from *Bacillus subtilis* Sfp, not all chimeric ACPs were initially obtained in the *holo* form. Samples that were identified as being a mixture of *apo*/*holo* by deconvolution of LCMS data using ESIprot^36^ were converted to the *holo* state via an on-column Sfp reaction during Ni-NTA purification step (**Table S10** and **Figures S1-S9**).^12,35^ Samples that lacked clear evidence of successful expression and purification (ACTα_2_-α_3_Ec, ACTα_3_-α_4_Ec, and ACTα_4_Ec) were not carried forward in the study. Notably, the observation that chimeras could be converted to their *holo*-form via the Sfp-catalyzed reaction suggests that the chimeric ACPs were able to bind and retain their function in the phosphopantetheinylation reaction.

### Analysis of ACT-backbone chimeras via SDS PAGE and circular dichroism (CD)

Despite both *holo*-AcpP and *holo*-ACT having a MW of ∼11 kDa (**Table S1** and **S2**, and **Figures S1** and **S2**), their migration patterns in SDS PAGE are markedly distinct. Under reducing conditions, *holo*-ACT migrates to its expected MW, whereas AcpP migrates to ∼20 kDa (**Figure S10**). This is consistent with previous observations by our lab and others, and has been attributed to the unusual molecular dimensions and/or charge distribution of the protein.^10,47^ Interestingly, all but one ACT-backbone chimeric ACP exhibited migration patterns similar to that of their template (ACT, C17S mutant). The ACT𝒩-α_2_Ec was the exception, migrating in a similar fashion to AcpP under both reducing and non-reducing conditions (**Figure S10**). These results indicate that the element of AcpP responsible for its unusually slow migration on SDS PAGE resides in loop 1 and helix II. Substitution of loop 1 or helix II alone did not significantly alter the migration. The overall global fold of the chimeric *holo*-ACPs were compared to that of AcpP and ACT *via* far-UV circular dichroism (CD). CD spectra of AcpP and ACT show strong negative features at 222 nm and 209 nm, which is consistent with previous studies and in alignment with the conserved primarily alpha helical structures of ACPs.^12^ The ACT-backbone chimeras maintained a similar alpha helical and random coil secondary structure as AcpP and ACT, although the percent helicity varied across chimeras (**Figure S11** and **Table S3**).

### Analysis of chimeric ACP binding to FabF via mechanism-based crosslinking assays

The *holo*-ACT-backbone chimeric ACPs were assessed for their ability to bind to FabF *via* a colorimetric crosslinking assay.^10^ In this assay, the terminal thiol of the ACP Ppant arm is converted into a mixed disulfide with 2-nitro-5-benzoate ion (TNB^−^) which thereby activates the site to form a selective, covalent crosslink with the active site cysteine of a cognate KS (Cys163 of FabF). Crosslinking can be monitored by the concomitant release of TNB^2–^, which absorbs at 412 nm and provides a visual and quantitative measure of the ACP-KS interaction (**Figure 2C**).

The extent of crosslinking of the ACT-backbone chimeric ACPs was initially monitored by eye, UV-Vis absorbance spectroscopy and SDS-PAGE. The following observations indicated potential crosslinking between the chimeric ACP and FabF: 1) a visual change in the reaction mixture from colorless to yellow, 2) the appearance of an absorbance band at 412 nm when visualized by UV-vis, 3) the presence of a higher molecular weight complex at ∼70 kDa consistent with the ACP-KS complex upon SDS PAGE analysis under non-reducing conditions, and 4) disappearance of the ∼10–20 kDa band corresponding to the stand-alone chimeric ACP-TNB^−^. The high concentrations of proteins used in this assay can lead to homodimer formation, which is important to consider when interpreting the significance of the crosslinking data. Using these methods, only two ACT-backbone chimeric ACPs were observed to crosslink to FabF: ACT𝒩-α_2_Ec and ACTα_2_Ec (**Figure S12**). For these two chimeras, the extent of crosslinking between the ACP and FabF was dependent on ACP concentration (**Figure 4**).

Size exclusion chromatography (SEC) was used to increase the rigor of the reaction analysis and breadth of undergraduate student training. In the case of ACT𝒩-α_2_Ec and ACTα_2_Ec, a higher molecular weight complex (eluting at 9 min, corresponding to the ACP-KS crosslink) and a small molecule product with λ_max_ at 412 nm (21 min, corresponding to TNB^2–^) were observed, consistent with the occurrence of a mechanism-based crosslink. These peaks were not observed for the other ACT-backbone chimeric ACPs. Instead, peaks at 11 min and 12–14 min were observed, corresponding to FabF and ACP-TNB^−^ respectively (**Figure S13**).

### Exchange of AcpP helix II for ACT helix II leads to loss of structure and FabF binding

To gain deeper insight into the role of helix II in AcpP recognition of partner enzymes, we assessed whether swapping helix II of AcpP for helix II of ACT would lead to a loss of functional interaction with FabF. Analysis of the mechanism-based colorimetric assay revealed that both Ec𝒩-α_2_ACT and Ecα_2_ACT chimeras did not form a mechanism-based crosslink with FabF (**Figure S14**). Investigation of the Ec𝒩-α_2_ACT and Ecα_2_ACT chimeras by CD suggested that an overall loss of secondary structure might be responsible for the lack of interaction between these two AcpP-backbone ACP chimeras and FabF (**Figure S15**). AcpP and ACT display helical structures that unfold upon increase in temperature, with observed T_m_s of 57.5 and 39.7 °C, respectively.^12^ We found that ACT𝒩-α_2_Ec displays a T_m_ of 30.9 °C whereas ACTα_2_Ec shows a T_m_ of 53.7 °C (**Figure S15, Table S4**), suggesting that ACTα_2_Ec maintains similar thermostability of the AcpP, whereas the Ec𝒩-α_2_ACT chimera behaves more similarly to the ACT. However, swapping either the AcpP helix II or the *N*-terminal sequence through helix II with the ACT region led to a significant loss in secondary structure (**Figure S15** and **Table S4**). Given that the helical nature of ACPs are a conserved structural feature that is thought to guide its functional interactions within the synthase,^7,48^ the loss of structure is likely a key driver for its loss of ability to bind to FabF.

### FabF-ACP binding analyses implicate helix II as a modular recognition element

Next, we explored the binding affinity of the chimeric ACPs in their *holo-*states via sedimentation velocity analytical ultracentrifugation (SV-AUC) and open surface plasmon resonance (OpenSPR) for several reasons. First, while our CD analyses indicate that installation of the TNB^−^ probe on the ACP Ppant arm does not perturb the secondary structure of the protein (**Figure S11**), it remains possible that the conversion of the Ppant arm free thiol to a mixed disulfide carrying bulky molecular cargo affects its interaction with binding partners. Second, both SV-AUC and OpenSPR provide insight into how the proteins interact in solution via non-covalent interactions and can provide further information about native binding affinities. Finally, these experiments enabled us to expand the pedagogical value of this project by increasing the breadth of biochemical concepts and techniques incorporated in the CURE.

SV-AUC analysis of ACTα_2_Ec and Ecα_2_ACT revealed that both proteins sediment as monomers under reducing conditions. Ecα_2_ACT was observed to sediment at a slower rate than ACTα_2_Ec, which is consistent with their respective migration patterns on SDS PAGE (**Figures S10** and **S16**). Binding analyses via SV-AUC and SPR revealed that ACTα_2_Ec binds to FabF with a weaker affinity than AcpP but stronger than ACT. In the case of SV-AUC, a change in the sedimentation boundary corresponding to weak binding^12^ was observed for FabF in the presence of ACTα_2_Ec but not Ecα_2_ACT (**Figure 5A&B** and **S16**). These results are consistent with the ACTα_2_Ec gaining the ability to bind to FabF and Ecα_2_ACT losing the ability to bind to FabF. A single SV-AUC titration experiment shows that the ACTα_2_Ec displays a K_D_ that is more than ten-fold higher than that of AcpP (∼140 μM vs ∼6 μM,^10^ respectively), which suggests the ACTα_2_Ec can bind to FabF albeit with a lower binding affinity (**Figure S16**). A similar trend in relative binding affinity was observed upon analysis of ACP-FabF binding by OpenSPR: ACTα_2_Ec bound to FabF with a K_D_∼seven-fold higher than AcpP (48 to 61 μM vs 8.5 to 14 μM), and ACT did not show evidence of interaction (**Figure 5C&D** and **Figure S17**). The observed attenuated binding of ACTα_2_Ec to FabF in these experiments was not apparent in the mechanism-based crosslinking experiments. This suggests the possibility that the installation of the TNB^−^ probe leads to a conformational change that strengthens the binding interaction between ACTα_2_Ec and FabF and/or might reflect limitations of assessing transient/weak interactions *via* covalent-trapping assays.

**Figure 5.**
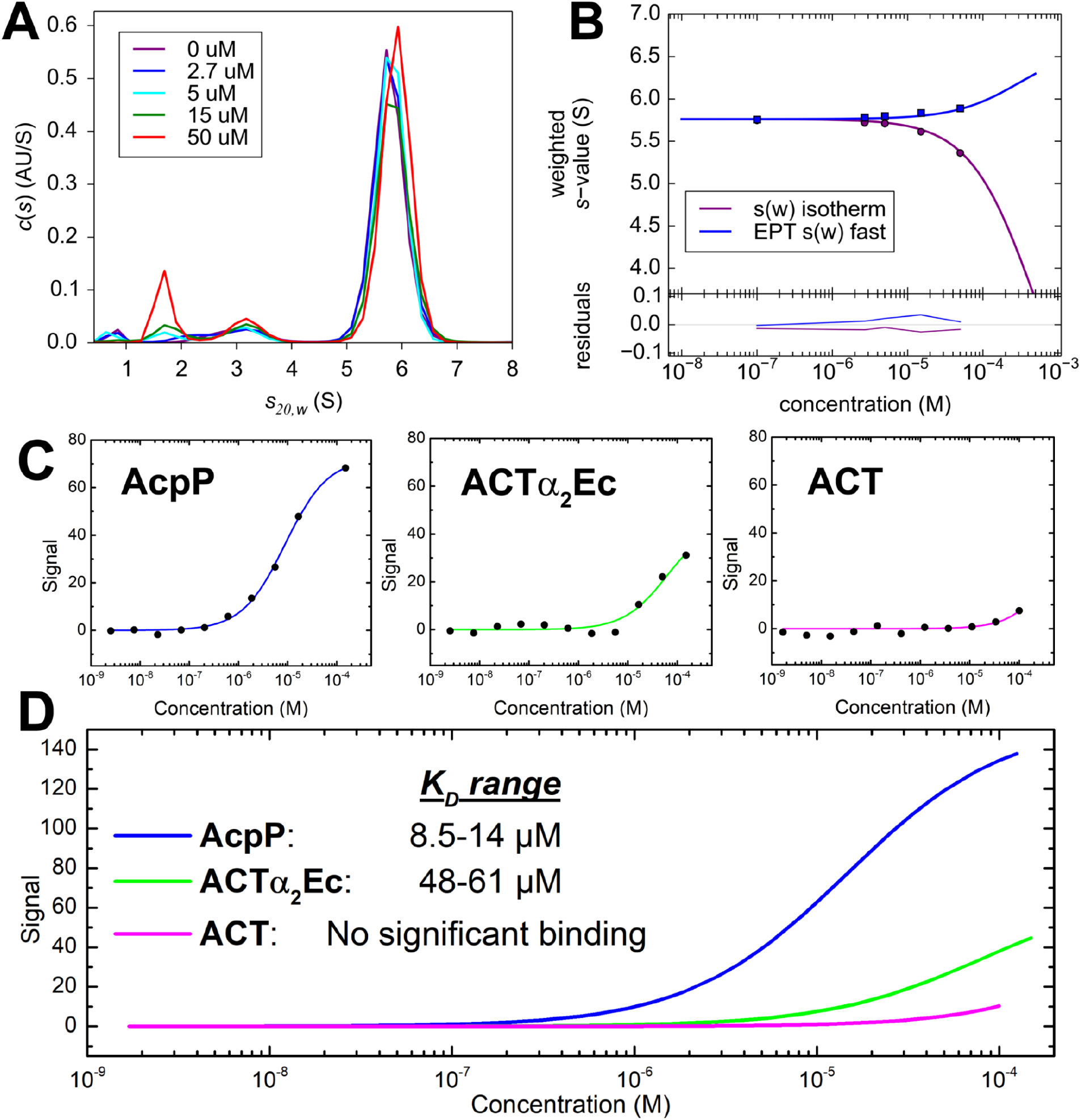
Dissociation constant studies of *holo*-ACTα_2_Ec. Dissociation constant (K_D_) for *holo*-ACTα_2_Ec (0–50 μM) interacting with FabF (17 μM) first determined from concentration-dependent sedimentation experiments (A-B). Both isotherms of weight-average sedimentation coefficient versus *holo*-ACTα_2_Ec concentration were fitted to A + B ↔ AB model with a K_D_ of 140 μM. The *holo*-ACTα_2_Ec peak is centered at 1.7 S. (A) Unnormalized c(s) distributions of FabF titrated with *holo*-ACTα_2_Ec. (B) EPT s(w) fast and s(w) weight-average sedimentation coefficient isotherms plotted against *holo*-ACTα_2_Ec concentrations. The fitted isotherms are shown as solid lines. The K_D_ of the three *holo*-ACPs (AcpP, ACTα_2_Ec, and ACT) were also studied by SPR analysis (C-D). (C) SPR result of *holo*-AcpP (left), *holo*-ACTα_2_Ec (center), and *holo*-ACT with FabF as a ligand. (D) Overlay of fitted triplicate SPR results of each ACP. The K_D_ range of *holo*-AcpP-FabF binding was observed as 8.5–15 μM and the K_D_ range of *holo*-ACTα_2_Ec -FabF was observed as 48–61 μM. No significant binding was observed between *holo*-ACT and FabF.

### Simulations support the role of helix II as a recognition site for ACP-FabF interactions

The interactions between ACPs and FabF were further analyzed using molecular dynamics (MD) simulations that started from related bound structures (see Supporting Information for details). First, a comparative analysis of the 1 μs simulations revealed that the root-mean-square deviation (RMSD) of the Ppant arm in the AcpP-FabF complex was relatively constant compared to that of the ACT-FabF complex (**Figure S18** and **Table S5**). These results are consistent with reduced stability for the non-cognate pair. Second, inter-protein salt bridges were analyzed using the Salt Bridges Plugin in Visual Molecular Dynamics (VMD), where a salt bridge was considered to have formed if the distance between an oxygen atom of an acidic residue and a nitrogen atom of a basic residue were within 3.2 Å for at least one frame of the simulation (**Tables S6**–**9**). Analysis of the two simulations revealed that the AcpP-FabF complex has nine pairs of residues that form a salt bridge for more than 10% of the trajectory, with an average percentage of 50% and low standard deviations in the residue-residue distances (**Figure 6**). The ACT-FabF complex only has six such pairs, with a lower average of 36% and higher standard deviations. Interestingly, eight of the nine salt bridges in the AcpP-FabF complex are formed between a helix II residue in the ACP and a FabF residue near the FabF active site. Three of the most significant salt bridges in the AcpP-FabF complex, Asp39-Lys65, Glu49-Lys128, and Asp39-Lys69, do not have homologs in the ACT-FabF complex. Perhaps the stability conferred by these residues, in addition to the stability granted by the Glu48-Arg127, Glu42-Lys69, and Asp36-Lys69 salt bridges, which do have homologs in the ACT-FabF complex, are essential for FabF to form a mechanistically relevant interaction with a particular ACP.

**Figure 6.**
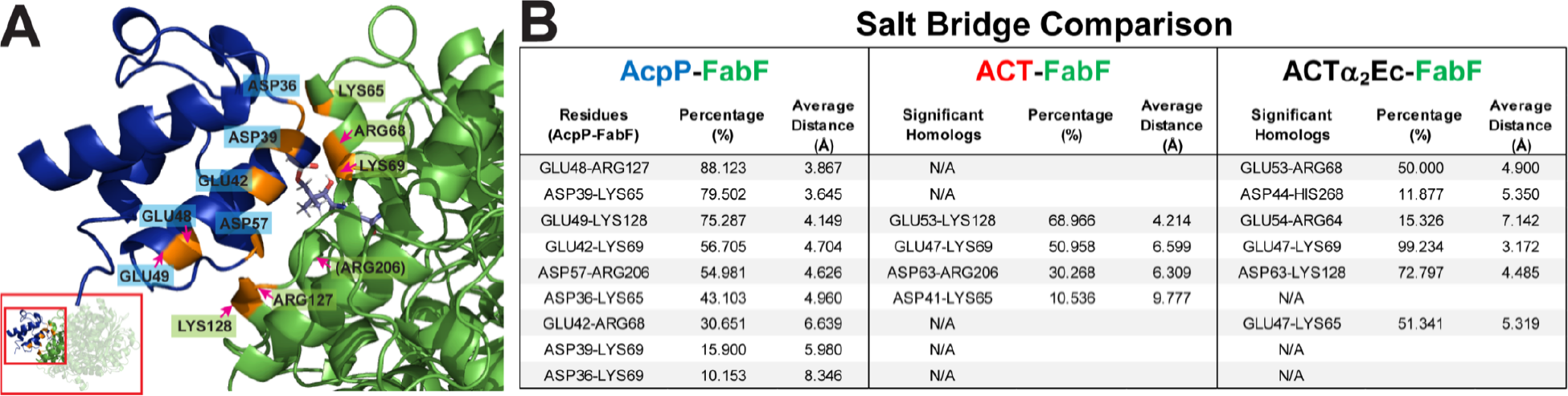
MD simulations point to specific salt bridges residing on helix II of AcpP that facilitate its interaction with FabF. (A) Simulated docking of AcpP (blue) and FabF (green) with amino acids engaging in salt bridges labeled and highlighted in orange. (B) List of AcpP, ACT, and ACTα_2_Ec amino acids that engage in salt bridges, along with the percent time that these were observed during the trajectory and the average distance of the interaction.

These findings align with observations from previous studies, including: 1) structural studies showing that AcpP helix II serves as a recognition site for the *E. coli* ketoreductase (KR, FabG),^27^ dehydratase (DH, FabA),^49^ acyl transferase (AT, FabD), and phosphopantetheinyl transferase (PPTase. AcpS);^50,51^ and 2) exchange of helix II of AcpP results in functional incompatibility *in vivo*.^26^ Moreover, these results are also in alignment with recent crystal structures of FabF and another *E. coli* elongating KS, FabB.^41,52^ Analysis of the FabB-AcpP and FabB-AcpP interfaces reveal electrostatic interactions between the negatively charged AcpP and positively charge KS. Specifically, AcpP residues Asp36, Asp39, Glu48, Glu49, Glu54, and Asp57 were found to make electrostatic interactions with two distinct positive patches on FabF. In addition, Leu38, Val41 and Met45 are involved in hydrophobic interactions with a different patch of FabF.^38^ Notably, eight of the eleven residues that are implicated in making connections between AcpP and FabF in the crystal structure reside in the helix II region of AcpP.

Additional preliminary molecular dynamics simulations starting from docking structures were performed in order to obtain the potential of mean force for pulling the ACP protein apart from the KS protein for five ACP structures: AcpP, ACT, Ecα_2_ACT, ACT𝒩*-*α_2_Ec, and ACTα_2_Ec (**Figure S19**). However, these simulations indicated that all five ACP structures would bind to KS.

This is in disagreement with the current experiments and suggests a need to more carefully explore the role of solution conditions and protein orientation in these free energy calculations.

## Conclusions

The molecular basis for the lack of interchangeability amongst ACPs across different synthases is an intriguing puzzle—its solution could lead to the strategic redesign of synthases capable of manufacturing diverse, structurally complex molecules. The observation that a type II PKS ACP (ACT) cannot substitute for AcpP in the *E. coli* type II FAS despite their conserved tertiary structures points to a fundamental question about what modular features of ACPs direct cross-synthase compatibility. Building off the premise that ACT does not interact with FabF whereas the cognate AcpP does, we designed, expressed, and purified chimeric ACPs with swapped secondary elements of ACT and AcpP. Next, we leveraged a suite of binding affinity assays and MD simulations to systematically identify the modular features of the AcpP that confer its functional interaction with its FabF. Our results implicate AcpP helix II as the recognition element in interactions with other biosynthetic partners. Our computational modeling of the AcpP-FabF interface shows that these electrostatic relations are maintained over time in the form of salt bridges. All but one of the significant salt bridges observed at the AcpP-FabF interface are formed between AcpP residues located on helix II and FabF residues near the KS active site. Taken together, these findings strengthen our mechanistic understanding of how ACPs interact with their biosynthetic partners. Future work will focus on exploring whether the chimeric ACPs retain their function to facilitate decarboxylative condensation and transacylation reactions with FabF, investigating their compatibility with other enzymatic partners (*e*.*g*. ketoreductases and other KSs), and connecting their binding affinity to their conformational dynamics (*e*.*g*. chain sequestration behavior). Together these studies will provide important insights into how ACPs can be engineered in the strategic redesign of synthases capable of producing novel chemical diversity.

## Methods

Additional experimental materials, methods, and results are provided in supporting information. Accession codes (UniProt) for proteins in this study are as follows: FabF KS: P0AAI5, AcpP: P0A6A8, ACT ACP: Q02054.

## Authors

All: Department of Chemistry, Haverford College, Haverford PA 19041, United States

## Author Contributions

C.L.A., L.K.C., Y.I.C., B.K. and A.S. Designed the project, conducted molecular biology and biochemical experiments, analyzed data, and contributed to writing the manuscript.

K.K.A., K.N.B., O.B., S. J. C., A. E. C-S., P. L. C-P., S. M. C., L.M.D., S.C.E., C. J. F., W. J. H., A. K., O.C.L., J.L., S. M. M., V. M., N. J. S. Helped design the project, conducted molecular biology and biochemical experiments and analyzed data.

C.E.W.C., G.S.H., S.H.M., C.A.D. and C.H.L. designed, conducted and analyzed computational simulations and helped write the manuscript.

All authors contributed to proofreading the manuscript.

## Supporting Information Available

This material is available free of charge *via* the Internet.

