## Supporting Information for "Engineered chimeras unveil swappable modular features of fatty acid and polyketide synthase acyl carrier proteins"

### Detailed Methods

**Design of Chimeric Acyl Carrier Proteins and Molecular Cloning**

Each pair of students designed and constructed two complementary ACP chimeras: one with an ACT-backbone and one AcpP-backbone. In total, the eight ACT-backbone and eight AcpP-backbone chimeras (**Figure 3**) were modeled using Phyre2[32] (**Table S1**) and constructed using Gibson Assembly (**Table S2**). Purchased gBlock DNA (Integrated DNA Technologies) was amplified *via* polymerase chain reaction (PCR) under the following conditions: 10 μL of 5x Q5 reaction buffer, 1 μL of 10 mM dNTP, 0.5 μL of Q5 DNA polymerase, 10 μL of Q5 GC enhancer, and 23.5 μL of nuclease free water for a total volume of 45 μL. The master mixture was then added to 2 μL of 10 ng/μL gBlock DNA, 2.5 μL of 10 μM forward primer, and 2.5 μL of 10 μM reverse primer to create a total volume of 52 μL. Using the thermocycler, the reaction mixture underwent a seven-step cycling protocol described as follows: 1. 98 °C 1 min, 2. 98 °C 30 sec, 3. 72 °C 30 sec, 4. 72 °C 30 sec, 5. Go to 2, 29 times, 6. 72 °C 10 min, and 7. 4 °C Hold). Successful amplification of the gBlock DNA was confirmed by DNA gel electrophoresis (1 % (w/v) agarose) and the amplified product was excised and purified using the Zymoclean DNA Clean & Concentrator Kit. The pET28a vector was linearized through digestion with NdeI and EcoRI for 4 hr at 37 °C and purified *via* DNA gel electrophoresis followed by gel extraction (Zymoclean).

Once the insert and vector were obtained as described above, each at a concentration of > 20 ng/μL after gel extraction, the two were combined in a 3:1 insert:vector molar ratio in a volume of 5 μL and added to 15 μL of Gibson Assembly® Master Mix (New England BioLabs), consisting of T5 Exonuclease, Phusion Polymerase, Taq Ligase, dNTPs, and $MgCl_2$ in Tris-HCl buffer. The resulting mixture was incubated for 1 hr at 50 °C and then transformed into DH5α competent cells *via* a 90-second, 42 °C heat shock. Resulting colonies were used to inoculate 10 mL LB/kan cultures (50 µg/mL) and grown overnight at 37 °C, 225 RPM. Plasmid DNA was isolated using a Qiagen QIAprep Spin Miniprep Kit and sequenced using standard T7 + T7term primers (Eurofins Genomics).

**Protein expression and purification**

AcpP, ACT,[10] and the chimeric ACPs were transformed into competent BAP1 *E. coli* cells which contained a phosphopantetheinyl transferase to convert the *apo* ACPs into their *holo* forms.[33] pXY-FabF, kanamycin-resistant, was transformed into *E. coli* BL21 cells. Cultures (800-mL) were inoculated with overnight seed culture (4-mL), incubated at 37 °C with shaking (200 rpm) until the OD (600 nm) 0.6-0.8 was reached. After induction with isopropyl β-D-1-thiogalactopyranoside (IPTG; 250 µM), the temperature was reduced to 18 °C for an additional 12–16 hours. Cells were harvested by centrifugation at 4,500 *g* for 15 minutes and resuspended in an ice-cold lysis buffer (50 mM sodium phosphate, pH 7.6, 300 mM NaCl, 10 mM imidazole, and 10% (v/v) glycerol) prior to lysis via sonication on ice using an XL-200 Microson sonicator (7 x 30 sec continuous pulse alternating with 30 sec rest, power level = 12 W). Cell debris were removed by centrifugation (17,000 *g* for 45 minutes). Supernatant was equilibrated with 3 mL Ni-NTA affinity resin (Gold Bio, pre-equilibrated with lysis buffer) for 1–2 hours at 4 °C. The protein-bound resin suspension was added to a poly-prep column (Bio-rad) and was allowed to settle before being washed with 50 mL of sodium phosphate buffer, pH 7.6, to bring the $A_{280}$ to ~0, indicating that any unbound protein was washed off the column.

Owing to *E. coli* harboring a native ACP phosphodiesterase, some of the expressed ACPs were isolated as *apo* and *holo* mixtures.[34] An additional, promiscuous, phosphopantetheinyl transferase Sfp-catalyzed reaction was used to convert the remaining *apo* ACPs into 100% *holo.*[35] Prior to applying the eluant onto a poly-prep (Bio Rad) column, 200 µL of expressed ACPs equilibrated resin was removed for analysis by LC-MS to evaluate the ratio of *apo*- vs *holo*-ACP in solution.[12] When necessary, ACPs were converted to 100% *holo* using an on-resin Sfp reaction, as described previously.[12] In brief, Ni-NTA resin harboring the $His_6$-tagged ACP was washed with lysis buffer until the $A_{280}$ reached baseline and then washed again using wash buffer until the $A_{280}$ once again reached zero. Resin was then mixed with 2 mL of an Sfp reaction mixture containing 50 mM sodium phosphate, pH 7.6, 10 mM $MgCl_2$, 2.5 mM coenzyme A (CoA), 15 mM fresh dithiothreitol (DTT), and 6 µM Sfp R4-4 phosphopantetheinyl transferase (a generous gift from Jun Lin's Lab at Georgia State University).[35] The Sfp-catalyzed addition of the Ppant arm onto remaining *apo*-ACP took place on column upon gentle rotation at room temperature. After reacting for 12–16 hrs, the resin was washed again with washing buffer until the excess coenzyme A was removed from the reaction and $A_{260}$ reached baseline. The Ni-bound *holo*-ACP sample was then eluted in 1-mL fractions with 10 mL of elution buffer (50 mM sodium phosphate, 100 mM NaCl, 300 mM imidazole, pH 7.6). Protein content was assessed by $A_{280}$ readings (see **Table S1** for extinction coefficients) using UV-Vis spectroscopy as well as Bradford assay. Full conversion of ACPs to their *holo* form was confirmed by LC-MS (see below). Fractions containing *holo*-ACPs were dialyzed against 50 mM sodium phosphate, pH 7.6 (3 K MWCO Slide-a-Lyzer Dialysis Cassettes; ThermoFisher), concentrated to 0.5–1 mM using Amicon Ultra-4 3K MWCO centrifugal filters, and stored at –80 °C for future use. FabF (KS) was purified as described previously.[10]

Fractions containing FabF (KS) were pooled together concentrated to about 400 μM using Amicon Ultra-4 10K MWCO centrifugal filters and stored in –80 °C with 10% (v/v) glycerol until future use. To prevent aggregation, stocks of FabF were dialyzed against 50 mM sodium phosphate pH 7.6 prior to each assay.

**Sodium Dodecyl Sulfate Polyacrylamide Gel Electrophoresis (SDS PAGE)**

SDS PAGE samples were prepared by mixing 2 μL of sample with either 2 μL of 6X loading dye and 8 μL of 50 μM sodium phosphate buffer for non-reducing samples and 6X loading dye mixed with freshly prepared 5% (v/v) beta-mercaptoethanol (BME) for reducing samples. 10 μL of each SDS PAGE sample was loaded in each gel well. Precision Plus Protein Standards (Bio-Rad, 10 μL) were loaded alongside the protein samples. SDS PAGE gels were run in 10X SDS running buffer (1% (w/v) SDS, 1.92 M glycine, 0.25 M Tris base, pH 8.3) at 120 V for ~60 min, or until the dye front had reached the bottom of the gel. The gels were washed with $ddH_2O$ at room temperature for 15 min, stained with GelCode Blue Stain Reagent for ~1 hr at room temperature with gentle shaking, and de-stained in $ddH_2O$ overnight at room temperature with gentle shaking prior to imaging.

**Molecular dynamics simulations.**

*ACP and KS Structural Model Acquisition:* The default settings for I-Tasser were selected, and the job was submitted to the server. The results were downloaded and analyzed using the outputs given in the index.html file.

*ACP-KS Docking:* The ACPs and FabF were each aligned with the experimental structure using the Alignment plugin in PyMOL with the “One to one” option and “align” method selected. The Mobile Selection was chosen to be either of the ACP or FabF, and the Target Selection was chosen to be the template structure. The number of cycles was set to 5 and the cutoff was 2.0.

*Conversion of* apo-*ACP to* holo-*ACP.* In Pymol, this conversion was accomplished *by* selecting the “Ppant_arm” object and clicking “copy to object” in the Action (A) toolbar, choosing the ACP of interest as the destination object. The docked ACP-FabF complex was saved by selecting the ACP and FabF and choosing File → Export Molecule…, then setting Selection to “enabled” and State to “-1 (current)” and selecting the “Write segment identifier (segi) column,” “Retain atom ids,” and “Write HEADER for every object” options. The resulting .pdb file was opened in a text editor and the serine residue of the ACP to which the Ppant arm would be attached was replaced with the “PNS” object found at the end of the file. This converted the PNS residue from an independent object to a residue within the ACP amino acid sequence, completing the process of converting the *apo*-ACP to *holo*-ACP, as well as completing the process of generating the ACP-KS-CLF structural model. The structure of the the non-native PNS residue was generated using bond lengths and approximate angles optimized in the MM2 forcefield, and validated with CHARMM MD parameters from CGenFF.[1]

*Preparation of Model for Molecular Dynamics*

The ACP-KS-CLF structural models were prepared for large-scale MD simulations using a local version of the GROMACS 2020.1 package and the “gromacs_prep.sh” script. Any water molecules present in the structure were removed. The complex was processed with the

CHARMM36 forcefield with the artificial residue PNS (which includes the covalently linked Ppant arm) with parameters derived from CGenFF.[1] The complex was centered in a cubic box with a distance of 1.0 nm from the complex to the edge of the box. The box was solvated with water (TIP3P) and neutralized with $Na^+$ or $Cl^-$ ions. The addition of ions and the energy minimization steps were completed with standard ions.mdp and minim.mdp files provided in a tutorial by Dr. Justin A. Lemkul from the Virginia Tech Department of Biochemistry (http://www.mdtutorials.com/gmx/lysozyme/index.html).

*Submission for Production Molecular Dynamics Simulation*

The prepared topology file and GROMACS structure file were transferred to Stampede2, a supercomputing cluster at the Texas Advanced Computing Center (TACC), for the MD production run. GROMACS 2018.3 was loaded and run in parallel on eight knl nodes with 64 MPI processes each (for a total of 512 MPI processes). 500 ps NVT and NPT equilibrations of the system were completed using standard nvt.mdp and npt.mdp settings files from the Lemkul tutorial. The 1 μs production simulations were run with 2 femtosecond time steps. Each simulation was completed over the course of thirteen sequential jobs, spanning about two total weeks of wall clock time.

*Analysis of Trajectory RMSD and SASA*

RMSD was evaluated in a Jupyter Notebook on the local machine using the MDAnalysis Python Library. The MD Analysis Universe was built with the MDAnalysis package before the RMSD analysis was used to evaluate the RMSD of the complex. The results were plotted using Matplotlib. The SASA of the Ppant arm was outputted using the gmx_sasa routine in Gromacs 2018.3.

*Analysis of Salt Bridges*

The existence of a salt bridge between a pair of amino acid residues was defined as an oxygen atom from one of the residues and a nitrogen atom from the other residue being within 3.2 Å of each other during at least one frame of the MD trajectory. The npt.gro file produced by the MD equilibration (the input GROMACS file for the production run) and a trimmed MD trajectory (~500 frames spanning the entire μs production run) were opened in Visual Molecular Dynamics 1.9.4 (VMD). The VMD Salt Bridge Analysis tool was used to evaluate each protein-protein complex (without solvent) for salt bridges, updating the selection for every frame. The resulting data was evaluated manually to categorize each salt bridge as an intra-KS link, an intra-ACP link, or an inter-protein link. After sorting, the percentage of time that the oxygen and nitrogen of a pair of residues were within 4 Å and the average distance of the oxygen-nitrogen pair were evaluated.

*Statistical Analysis of RMSD, SASA, and Salt Bridge Results*

The statistical differences between the RMSD, SASA, and salt bridge data sets of different groups of ACP-KS pairs were evaluated using a two-tailed Student's t-test. In all cases, the null hypothesis was that there was no significant difference between the data sets being compared. Additionally, the alternative hypothesis was that the data sets were significantly different from each other. The standard deviation of Group N, $s_n$, was determined using the following formula:

$$s_n = \sqrt{\sum_{i=0}^{N} \frac{(x_i - \mu)^2}{(N-1)}}$$

where $x_i$ is the value of an individual data point in the group, $\mu$ is the sample mean for the group, and N is the number of samples in the group. The standard error, $s_d$, between two groups was determined using the following formula:

$$s_d = \sqrt{\frac{s_1}{N_1} + \frac{s_2}{N_2}}$$

where $s_1$ is the standard deviation of Group 1, $s_2$ is the standard deviation of Group 2, $N_1$ is the sample size of Group 1, and $N_2$ is the sample size of Group 2. The t-score, t, was calculated using the following formula:

$$t = \frac{|\mu_1 - \mu_2|}{s_d}$$

where $\mu_1$ is the sample mean for Group 1, $\mu_2$ is the sample mean for Group 2, and $s_d$ is the standard error between the two groups. The degrees of freedom, df, were determined by adding the number of samples in each group and subtracting two:

$$df = N_1 + N_2 - 2$$

where $N_1$ and $N_2$ are the number of samples in Group 1 and Group 2, respectively. Finally, the calculated values for *t* and *df* were used with a standard t-table to determine a p-value for the comparison of the pair of data sets. A p-value less than 0.05, corresponding to a less than 5% probability of the data being a result of random chance, was chosen as the cutoff for statistical significance.

**Free energy calculations.**

Free energy calculations were performed using the python based toolkits OpenMM, mdtraj, and pymbar.[1–3] Bound KS+ACP structures were obtained from the previously completed docking analysis for five ACP structures: AcpP, ACT, Ec$\alpha_2$ACT, ACT$\mathcal{N}$-$\alpha_2$Ec, and ACT$\alpha_2$Ec. Following processing to add missing hydrogens in CHARMM-GUI, each structure was described using the CHARMM36 force field and solvated in TIP3P water and NaCl counterions to neutrality.[5–7] CGenFF parameters were used for the Ppant arm, as described in the Materials and Methods section. A 1000 kcal/mol/Å$^2$ harmonic force was applied to the backbone atoms (C, CA, and N), holding them in place. Then the systems were each subjected to the same equilibration procedure: energy minimization, randomization of velocities to 5 K, slowly increasing the temperature to 300 K over 1 ns under NPT (1 atm) conditions, slowly removing the backbone constraint over 1 ns under NPT (300 K, 1 atm) conditions, 1 ns of NPT (300 K, 1 atm) simulation with no constraints, and finally 2 ns of NVT (300 K) simulation. Langevin dynamics were used to hold temperature constant and a Monte Carlo barostat was used to control pressure.[8]

Following equilibration, the systems were prepared for umbrella sampling. 10 systems were prepared where the center of mass distance between the ACP and KS was held still with a 0.1 kcal/mol/Å$^2$ harmonic force. In each simulation, the complex was held to a different distance between 30 and 60 Å, evenly spaced between the 10 systems. After 2 ns of NVT (300 K) equilibration, the systems were run in NVT for 60 ns, collecting samples every 300 ps. This low sampling frequency was chosen to minimize time correlation between samples. After these simulations were complete, the center of mass distance between the KS and the ACP was calculated for every sample and the multistate Bennett acceptance ratio method was used to calculate the free energy differences between the bound states.[3] These free energy differences were then used to calculate a potential of mean force along the center of mass distance coordinate.[4,9]

These potentials of mean force are effectively free energy surfaces for a simple one dimensional coordinate,[10] and are shown in Figure S19. Values are shifted so that the unbound state, where the proteins are interacting only with solvent, are at zero for each ACP protein. All potentials of mean force imply a favorable interaction between the ACP and KS, since the free energy decreases as the proteins are brought together. This is in disagreement with the current experimental observations, where some of the ACPs are seen to bind to the KS but others do not. In our future simulations we will investigate the source of this discrepancy. Preliminary analyses suggest that the origin might have to do with the solution conditions, protein orientational freedom, or the impact of the two KS domains.

#### Tables and Figures

**Table S1.** Extinction coefficients, molecular weights, and PyMOL models of ACPs used in this study. PyMOL figures for ACP chimeras were generated using Phyre2 software.

| **Protein** | **MW (Da)** | **Extinction Coefficient ($M^{-1}cm^{-1}$)** | **PyMOL Model** |
|---|---|---|---|
| AcpP<br><br>(Model adapted from PDB: 6OLT) | 10802.84 | 1490 | 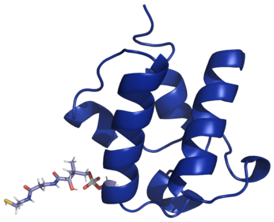 |
| ACT<br>(C17S mutant, PDB: 2K0X) | 11395.55 | 2980 | 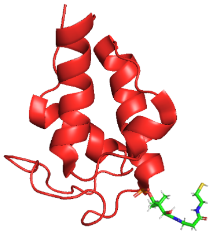 |
| FabF<br>(PDB: 2GFW) | 44231.01 | 25900 | 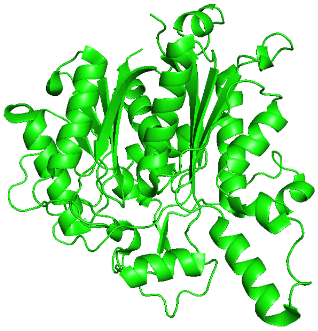 |
| **Chimeric ACPs with ACT-backbone** | | | |

| **Protein** | **MW (Da)** | **Extinction Coefficient ($M^{-1}cm^{-1}$)** | **PyMOL Model** |
|---|---|---|---|
| ACT$\mathcal{N}$-$\alpha_1$Ec | 11293.5 | 2980 | 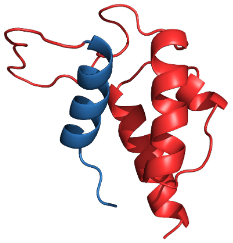 |
| ACT$\mathcal{N}$-$\alpha_2$Ec | 10937.1 | NA | 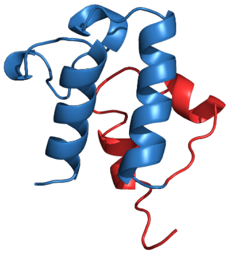 |
| ACT$\alpha_2$Ec | 11409.5 | 1490 | 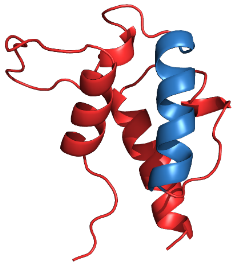 |
| ACT$\alpha_2$-$\alpha_3$Ec | 11555.6 | 1490 | 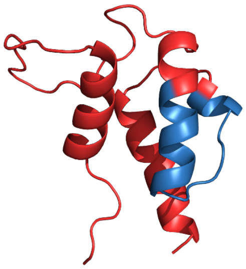 |
| ACT$l_2$Ec | 11497.59 | 2980 | 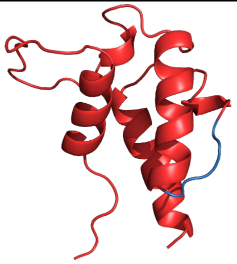 |
| ACT$l_2$-$l_3$Ec | 11585.70 | 2980 | 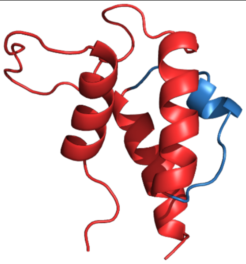 |

| | | | |
|---|---|---|---|
| ACT$\alpha_3$-$\alpha_4$Ec | 11473.57 | 4470 | 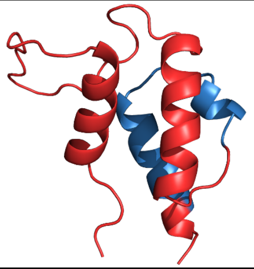 |
| ACT$\alpha_4$Ec | 11129.17 | 4470 | 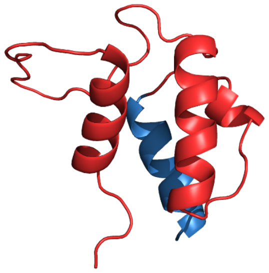 |
| **Chimeric ACPs with AcpP(“Ec”)-backbone** | | | |
| **Protein** | **MW (Da)** | **Extinction Coefficient ($M^{-1}cm^{-1}$)** | **PyMOL Model** |
| Ec$\mathcal{N}$-$\alpha_1$ACT | 10904.89 | 1490 | 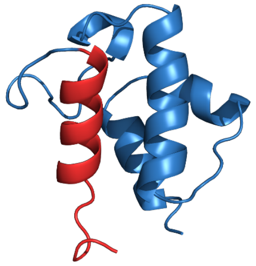 |
| Ec$\mathcal{N}$-$\alpha_2$ACT | 11261.28 | 4470 | 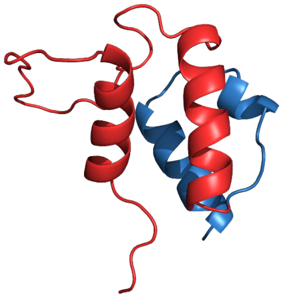 |
| Ec$\alpha_2$ACT | 10788.87 | 2980 | 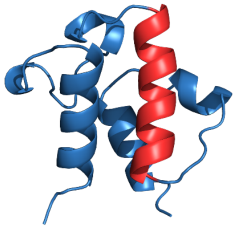 |

| | | | |
|---|---|---|---|
| Ec$\alpha_2$-$\alpha_3$ACT | 10641.83 | 2980 | 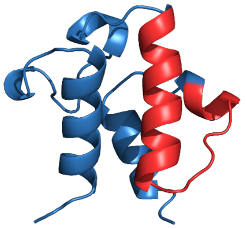 |
| Ec$l_2$ACT | 10700.80 | 1490 | 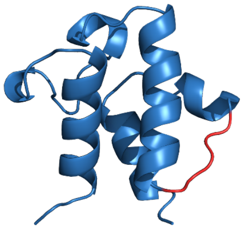 |
| Ec$l_2$-$l_3$ACT | 10612.69 | 1490 | 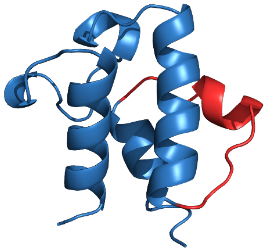 |
| Ec$\alpha_3$-$\alpha_4$ACT | 10724.82 | NA | 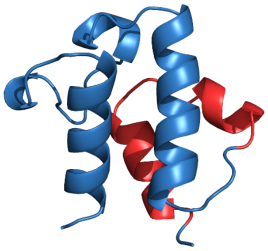 |
| Ec$\alpha_4$ACT | 11069.22 | NA | 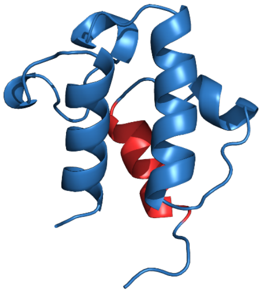 |

**Table S2.** Primer, gBlock insert, and amino acid sequences. The nucleotide sequences of the primers used for the amplification of the gBlock DNA and of the gBlock that was inserted into a pET28a vector to create novel plasmids. The amino acid sequences of the wild type ACPs, FabF, and the chimeric ACPs encoded by the newly assembled plasmids. MGSSHHHHHHSSGLVPRGSH = pET28a leader sequence including the $His_6$-tag used for affinity column purification.

| **Protein** | **Amino Acid Sequence and for ACP chimeras:<br>Forward Primer, Reverse Primer, and gBlock Insert** |
|---|---|
| AcpP | MGSSHHHHHHSSGLVPRGSHMSTIEERVKKIIGEQLGVKQEEVTN<br>NASFVEDLGADSLDTVELVMALEEEFDTEIPDEEAEKITTVQAAID<br>YINGHQA |
| ACT<br>(C17S mutant) | MGSSHHHHHHSSGLVPRGSHMATLLTTDDLRRALVESAGETDGT<br>DLSGDFLDLRFEDIGYDSLALMETAARLESRYGVSIPDDVAGRVD<br>TPRELLDLINGALAEAA |
| FabF | MRGSHHHHHHGSACVSKRRVVVTGLGMLSPVGNTVESTWKALL<br>AGQSGISLIDHFDTSAYATKFAGLVKDFNCEDIISRK<br>EQRKMDAFIQYGIVAGVQAMQDSGLEITEENATRIGAAIGSGIGG<br>LGLIEENHTSLMNGGPRKISPFFVPSTIVNMVAGH<br>LTIMYGLRGPSISIATACTSGVHNIGHAARIIAYGDADVMVAGGA<br>EKASTPLGVGGFGAARALSTRNDNPQAASRPWDKE<br>RDGFVLGDGAGMLVLEEYEHAKKRGAKIYAELVGFGMSSDAYH<br>MTSPPENGAGAALAMANALRDAGIEASQIGYVNAHGT<br>STPAGDKAEAQAVKTIFGEAASRVLVSSTKSMTGHLLGAAGAVE<br>SIYSILALRDQAVPPTINLDNPDEGCDLDFVPHEAR<br>QVSGMEYTLCNSFGFGGTNGSLIFKKI |
| **Chimeric ACPs with ACT-backbone** | |
| ACT$\mathcal{N}$-$\alpha_1$Ec | Forward Primer: gtgccgcgcggcagccatatgGCAACCCTGCTGACC<br>Reverse Primer: gtcgacggagctcgaattcTCATGCCGCCTCGGC<br>gBlock Insert:<br>AAAAAACATATGAGCACTATCGAAGAACGCGTTAAGAAAATT<br>ATCGGCGAACAGCTGGGTGAGACGGACGGGACGGACCTGTCC<br>GGCGACTTCCTCGACCTCCGCTTCGAGGACATCGGGTACGACT<br>CGCTCGCCCTGATGGAGACCGCGGCGCGACTCGAGAGCCGGT<br>ACGGCGTGTCCATACCCGACGACGTCGCCGGCCGCGTCGACAC<br>GCCGCGAGAGCTGCTCGACCTGATCAACGGCGCACTGGCCGA<br>GGCGGCATGAGAATTCAAAAAA<br>Amino Acid Sequence:<br>MGSSHHHHHHSSGLVPRGSHMSTIEERVKKIIGEQLGETDGTDLS<br>GDFLDLRFEDIGYDSLALMETAARLESRYGVSIPDDVAGRVDTPR<br>ELLDLINGALAEAA |
| ACT$\mathcal{N}$-$\alpha_2$Ec | Forward Primer: gtgccgcgcggcagccatatgGCAACCCTGCTGACC<br>Reverse Primer: gtcgacggagctcgaattcTCATGCCGCCTCGGC<br>gBlock Insert:<br>AAAAAACATATGAGCACTATCGAAGAACGCGTTAAGAAAATT<br>ATCGGCGAACAGCTGGGCGTTAAGCAGGAAGAAGTTACCAAC |

| | |
|---|---|
| | AATGCTTCTTTCGTTGAAGACCTGGGCGCGGATTCTCTTGACAC CGTTGAGCTGGTAATGGCTCTGGAAGAAGAGTTTGATGTGTCC ATACCCGACGACGTCGCCGGCCGCGTCGACACGCCGCGAGAG CTGCTCGACCTGATCAACGGCGCACTGGCCGAGGCGGCATGAG AATTCAAAAAA<br>Amino Acid Sequence:<br>MGSSHHHHHHSSGLVPRGSHMSTIEERVKKIIGEQLGVKQEEVTN NASFVEDLGADSLDTVELVMALEEEFDVSIPDDVAGRVDTPRELL DLINGALAEAA |
| ACTα$_2$Ec | Forward Primer: gtgccgcgcggcagccatatgGCAACCCTGCTGACC<br>Reverse Primer: gtcgacggagctcgaattcTCATGCCGCCTCGGC<br>gBlock Insert:<br>AAAAAACATATGGCAACCCTGCTGACCACCGACGATCTGCGCC GCGCCCTCGTGGAGAGCGCCGGTGAGACGGACGGGACGGACC TGTCCGGCGACTTCCTCGACCTCCGCTTCGAGGACATCGGGTA CGACTCGCTTGACACCGTTGAGCTGGTAATGGCTCTGGAAGAA GAGTTTGGCGTGTCCATACCCGACGACGTCGCCGGCCGCGTCG ACACGCCGCGAGAGCTGCTCGACCTGATCAACGGCGCACTGGC CGAGGCGGCATGAGAATTCAAAAA<br>Amino Acid Sequence:<br>MGSSHHHHHHSSGLVPRGSHMATLLTTDDLRRALVESAGETDGT DLSGDFLDLRFEDIGYDSLDTVELVMALEEEFGVSIPDDVAGRVD TPRELLDLINGALAEAA |
| ACTα$_2$-α$_3$Ec | Forward Primer: gtgccgcgcggcagccatatgGCAACCCTGCTGACC<br>Reverse Primer: gtcgacggagctcgaattcTCATGCCGCCTCGGC<br>gBlock Insert:<br>AAAAAACATATGGCAACCCTGCTGACCACCGACGATCTGCGCC GCGCCCTCGTGGAGAGCGCCGGTGAGACGGACGGGACGGACC TGTCCGGCGACTTCCTCGACCTCCGCTTCGAGGACATCGGGTA CGACTCGCTGGATACCGTGGAACTGGTGATGGCGCTGGAAGA AGAATTTGATACCGAAATTCCGGATGAAGAAGCGGGCCGCGT CGACACGCCGCGAGAGCTGCTCGACCTGATCAACGGCGCACTG GCCGAGGCGGCATGAGAATTCAAAAAA<br>Amino Acid Sequence:<br>MGSSHHHHHHSSGLVPRGSHMATLLTTDDLRRALVESAGETDGT DLSGDFLDLRFEDIGYDSLDTVELVMALEEEFDTEIPDEEAGRVDT PRELLDLINGALAEAA |
| ACT*l*$_2$Ec | Forward Primer: gtgccgcgcggcagccatatgGCAACCCTGCTGACC<br>Reverse Primer: gtcgacggagctcgaattcTCATGCCGCCTCGGC<br>gBlock Insert:<br>AAAAAACATATGGCAACCCTGCTGACCACCGACGATCTGCGCC GCGCCCTCGTGGAGAGCGCCGGTGAGACGGACGGGACGGACC TGTCCGGCGACTTCCTCGACCTCCGCTTCGAGGACATCGGGTA CGACTCGCTCGCCCTGATGGAGACCGCGGCGCGACTCGAGAGC CGGTACGATACTGAGATTCCGGACGACGTCGCCGGCCGCGTCG |

| | |
|---|---|
| | ACACGCCGCGAGAGCTGCTCGACCTGATCAACGGCGCACTGGC CGAGGCGGCATGAGAATTCAAAAAA<br>Amino Acid Sequence:<br>MGSSHHHHHHSSGLVPRGSHMATLLTTDDLRRALVESAGETDGT DLSGDFLDLRFEDIGYDSLALMETAARLESRYDTEIPDDVAGRVD TPRELLDLINGALAEAA |
| ACT*l*$_2$-*l*$_3$Ec | Forward Primer: gtgccgcgcggcagccatatgGCAACCCTGCTGACC<br>Reverse Primer: gtcgacggagctcgaattcTCATGCCGCCTCGGC<br>gBlock Insert:<br>AAAAAACATATGGCAACCCTGCTGACCACCGACGATCTGCGCC GCGCCCTCGTGGAGAGCGCCGGTGAGACGGACGGGACGGACC TGTCCGGCGACTTCCTCGACCTCCGCTTCGAGGACATCGGGTA CGACTCGCTCGCCCTGATGGAGACCGCGGCGCGACTCGAGAGC CGGTACGATACTGAGATTCCGGACGAAGAAGCTGAGAAAATC ACCACCCCGCGAGAGCTGCTCGACCTGATCAACGGCGCACTGG CCGAGGCGGCATGAGAATTCAAAAAA<br>Amino Acid Sequence:<br>MGSSHHHHHHSSGLVPRGSHMATLLTTDDLRRALVESAGETDGT DLSGDFLDLRFEDIGYDSLALMETAARLESRYDTEIPDEEAEKITT PRELLDLINGALAEAA |
| ACT$\alpha_3$-$\alpha_4$Ec | Forward Primer: gtgccgcgcggcagccatatgATGGCAACCCTGCTG<br>Reverse Primer: gtcgacggagctcgaattcTCATGCCGCCTCGGC<br>gBlock Insert:<br>AAAAAACATATGGCAACCCTGCTGACCACCGACGATCTGCGCC GCGCCCTCGTGGAGAGCGCCGGTGAGACGGACGGGACGGACC TGTCCGGCGACTTCCTCGACCTCCGCTTCGAGGACATCGGGTA CGACTCGCTCGCCCTGATGGAGACCGCGGCGCGACTCGAGAGC CGGTACGGCGTGTCCATACCGGACGAAGAAGCTGAGAAAATC ACCACCGTTCAGGCTGCCATTGATTACATCAACGGCCACCTGG CCGAGGCGGCATGAGAATTCAAAAAA<br>Amino Acid Sequence:<br>MGSSHHHHHHSSGLVPRGSHMATLLTTDDLRRALVESAGETDGT DLSGDFLDLRFEDIGYDSLALMETAARLESRYGVSIPDEEAEKITT VQAAIDYINGHLAEAA |
| ACT$\alpha_4$Ec | Forward Primer: gtgccgcgcggcagccatatgGCAACCCTGCTGACC<br>Reverse Primer: gtcgacggagctcgaattcTCATGCCGCCTCGGC<br>gBlock Insert:<br>AAAAAACATATGATGGCAACCCTGCTGACCACCGACGATCTGC GCCGCGCCCTCGTGGAGAGCGCCGGTGAGACGGACGGGACGG ACCTGTCCGGCGACTTCCTCGACCTCCGCTTCGAGGACATCGG GTACGACTCGCTCGCCCTGATGGAGACCGCGGCGCGACTCGAG AGCCGGTACGGCGTGTCCATACCCGACGACGTCGCCGGCCGCG TCGACACGGTTCAGGCTGCCATTGATTACATCAACGGCCACCA GGCGTAAGAATTCAAAAAA<br>Amino Acid Sequence: |

| | |
|---|---|
| | MGSSHHHHHHSSGLVPRGSHMATLLTTDDLRRALVESAGETDGT DLSGDFLDLRFEDIGYDSLALMETAARLESRYGVSIPDDVAGRVD TVQAAIDYINGHQA |
| **Chimeric ACPs with AcpP("Ec")-backbone** | |
| Ec$\mathcal{N}$-$\alpha_1$ACT | Forward Primer: gtgccgcgcggcagccatatgAGCACTATCGAA<br>Reverse Primer: gtcgacggagctcgaattcTTACGCCTGGTGGCC<br>gBlock Insert:<br>AAAAAACATATGGCAACCCTGCTGACCACCGACGATCTGCGCC GCGCCCTCGTGGAGAGCGCCGGCGTTAAGCAGGAAGAAGTTA CCAACAATGCTTCTTTCGTTGAAGACCTGGGCGCGGATTCTCTT GACACCGTTGAGCTGGTAATGGCTCTGGAAGAAGAGTTTGATA CTGAGATTCCGGACGAAGAAGCTGAGAAAATCACCACCGTTC AGGCTGCCATTGATTACATCAACGGCCACCAGGCGTAAGAATT CAAAAAA<br>Amino Acid Sequence:<br>MGSSHHHHHHSSGLVPRGSHMATLLTTDDLRRALVESAGVKQEE VTNNASFVEDLGADSLDTVELVMALEEEFDTEIPDEEAEKITTVQ AAIDYINGHQA |
| Ec$\mathcal{N}$-$\alpha_2$ACT | Forward Primer: gtgccgcgcggcagccatatgAGCACTATCGAA<br>Reverse Primer: gtcgacggagctcgaattcTTACGCCTGGTGGCC<br>gBlock Insert:<br>AAAAAACATATGGCAACCCTGCTGACCACCGACGATCTGCGCC GCGCCCTCGTGGAGAGCGCCGGTGAGACGGACGGGACGGACC TGTCCGGCGACTTCCTCGACCTCCGCTTCGAGGACATCGGGTA CGACTCGCTCGCCCTGATGGAGACCGCGGCGCGACTCGAGAGC CGGTACGGCACTGAGATTCCGGACGAAGAAGCTGAGAAAATC ACCACCGTTCAGGCTGCCATTGATTACATCAACGGCCACCAGG CGTAAGAATTCAAAAAA<br>Amino Acid Sequence:<br>MGSSHHHHHHSSGLVPRGSHMATLLTTDDLRRALVESAGETDGT DLSGDFLDLRFEDIGYDSLALMETAARLESRYGTEIPDEEAEKITT VQAAIDYINGHQA |
| Ec$\alpha_2$ACT | Forward Primer: gtgccgcgcggcagccatatgAGCACTATCGAA<br>Reverse Primer: gtcgacggagctcgaattcTTACGCCTGGTGGCC<br>gBlock Insert:<br>AAAAAACATATGAGCACTATCGAAGAACGCGTTAAGAAAATT ATCGGCGAACAGCTGGGCGTTAAGCAGGAAGAAGTTACCAAC AATGCTTCTTTCGTTGAAGACCTGGGCGCGGATTCTCTCGCCCT GATGGAGACCGCGGCGCGACTCGAGAGCCGGTACGATACTGA GATTCCGGACGAAGAAGCTGAGAAAATCACCACCGTTCAGGC TGCCATTGATTACATCAACGGCCACCAGGCGTAAGAATTCAAA AAA<br>Amino Acid Sequence:<br>MGSSHHHHHHSSGLVPRGSHMSTIEERVKKIIGEQLGVKQEEVTN NASFVEDLGADSLALMETAARLESRYDTEIPDEEAEKITTVQAAID YINGHQA |

| | |
|---|---|
| Ec$\alpha_2$-$\alpha_3$ACT | Forward Primer: gtgccgcgcggcagccatatgAGCACTATCGAA<br>Reverse Primer: gtcgacggagctcgaattcTTACGCCTGGTGGCC<br>gBlock Insert:<br>AAAAAACATATGAGCACTATCGAAGAACGCGTTAAGAAAATT<br>ATCGGCCAACAGCTGGGCGTTAAGCAGGAAGAAGTTACCAAC<br>AATGCTTCTTTCGTTGAAGACCTGGGCGCGGATTCTCTCGCCCT<br>GATGGAGACCGCGGCGCGACTCGAGAGCCGGTACGGCGTGTC<br>CATACCCGACGACGTCGCCGAGAAAATCACCACCGTTCAGGCT<br>GCCATTGATTACATCAACGGCCACCAGGCGTAAGAATTCAAAA<br>AA<br>Amino Acid Sequence:<br>MGSSHHHHHHSSGLVPRGSHMSTIEERVKKIIGQQLGVKQEEVTN<br>NASFVEDLGADSLALMETAARLESRYGVSIPDDVAEKITTVQAAI<br>DYINGHQA |
| Ec$l_2$ACT | Forward Primer: gtgccgcgcggcagccatatgAGCACTATCGAA<br>Reverse Primer: gtcgacggagctcgaattcTTACGCCTGGTGGCC<br>gBlock Insert:<br>AAAAAACATATGAGCACTATCGAAGAACGCGTTAAGAAAATT<br>ATCGGCGAACAGCTGGGCGTTAAGCAGGAAGAAGTTACCAAC<br>AATGCTTCTTTCGTTGAAGACCTGGGCGCGGATTCTCTTGACAC<br>CGTTGAGCTGGTAATGGCTCTGGAAGAAGAGTTTGGCGTGTCC<br>ATACCCGACGAAGAAGCTGAGAAAATCACCACCGTTCAGGCT<br>GCCATTGATTACATCAACGGCCACCAGGCGTAAGAATTCAAAA<br>AA<br>Amino Acid Sequence:<br>MGSSHHHHHHSSGLVPRGSHMSTIEERVKKIIGEQLGVKQEEVTN<br>NASFVEDLGADSLDTVELVMALEEEFGVSIPDEEAEKITTVQAAID<br>YINGHQA |
| Ec$l_2$-$l_3$ACT | Forward Primer: gtgccgcgcggcagccatatgAGCACTATCGAA<br>Reverse Primer: gtcgacggagctcgaattcTTACGCCTGGTGGCC<br>gBlock Insert:<br>AAAAAACATATGAGCACTATCGAAGAACGCGTTAAGAAAATT<br>ATCGGCGAACAGCTGGGCGTTAAGCAGGAAGAAGTTACCAAC<br>AATGCTTCTTTCGTTGAAGACCTGGGCGCGGATTCTCTTGACAC<br>CGTTGAGCTGGTAATGGCTCTGGAAGAAGAGTTTGGCGTGTCC<br>ATACCCGACGACGTCGCCGGCCGCGTCGACACGGTTCAGGCTG<br>CCATTGATTACATCAACGGCCACCAGGCGTAAGAATTCAAAAA<br>A<br>Amino Acid Sequence:<br>MGSSHHHHHHSSGLVPRGSHMSTIEERVKKIIGEQLGVKQEEVTN<br>NASFVEDLGADSLDTVELVMALEEEFGVSIPDDVAGRVDTVQAAI<br>DYINGHQA |
| Ec$\alpha_3$-$\alpha_4$ACT | Forward Primer: gtgccgcgcggcagccatatgATGAGCACTATCGAA<br>Reverse Primer: gtcgacggagctcgaattcTCATTACGCCTGTGC<br>gBlock Insert: |

| | |
|---|---|
| | AAAAAACATATGAGCACTATCGAAGAACGCGTTAAGAAAATT ATCGGCGAACAGCTGGGCGTTAAGCAGGAAGAAGTTACCAAC AATGCTTCTTTCGTTGAAGACCTGGGCGCGGATTCTCTTGACAC CGTTGAGCTGGTAATGGCTCTGGAAGAAGAGTTTGATACTGAG ATTCCCGACGACGTCGCCGGCCGCGTCGACACGCCGCGAGAGC TGCTCGACCTGATCAACGGCGCACAGGCGTAATGAGAATTCAA AAAA<br>Amino Acid Sequence:<br>MGSSHHHHHHSSGLVPRGSHMSTIEERVKKIIGEQLGVKQEEVTN NASFVEDLGADSLDTVELVMALEEEFDTEIPDDVAGRVDTPRELL DLINGAQA |
| Ecα$_4$ACT | Forward Primer: gtgccgcgcggcagccatatgAGCACTATCGAA<br>Reverse Primer: gtcgacggagctcgaattcTTACGCCTGGTGGCC<br>gBlock Insert:<br>AAAAAACATATGAGCACTATCGAAGAACGCGTTAAGAAAATT ATCGGCGAACAGCTGGGCGTTAAGCAGGAAGAAGTTACCAAC AATGCTTCTTTCGTTGAAGACCTGGGCGCGGATTCTCTTGACAC CGTTGAGCTGGTAATGGCTCTGGAAGAAGAGTTTGATACTGAG ATTCCGGACGAAGAAGCTGAGAAAATCACCACCCCGCGAGAG CTGCTCGACCTGATCAACGGCGCACTGGCCGAGGCGGCATGAG AATTCAAAAAA<br>Amino Acid Sequence:<br>MGSSHHHHHHSSGLVPRGSHMSTIEERVKKIIGEQLGVKQEEVTN NASFVEDLGADSLDTVELVMALEEEFDTEIPDEEAEKITTPRELLD LINGALAEAA |

**Table S3.** Calculated the 222/208 absorbance ratio of *holo* AcpP, ACT, and ACT-backbone ACP chimeras from CD spectra.

| ACP | 222/208 |
|---|---|
| AcpP | 0.84 |
| ACT | 0.86 |
| ACT$\mathcal{N}$-$\alpha_1$Ec | 0.76 |
| ACT$\mathcal{N}$-$\alpha_2$Ec | 0.87 |
| ACT$\alpha_2$Ec | 0.85 |
| ACT$\alpha_2$-$\alpha_3$Ec | 1.68 |
| ACT$l_2$Ec | 0.66 |
| ACT$l_2$-$l_3$Ec | 0.64 |
| ACT$\alpha_3$-$\alpha_4$Ec | 0.72 |
| ACT$\alpha_4$Ec | 0.51 |

**Table S4.** Thermostabilities, and changes in enthalpy and entropy of unfolding of select ACPs. AcpP and ACT data from Rivas 2018[11] whereas other data is from the current work.

| Protein | $T_m$ (°C) | $\Delta H_m$ (kcal/mol) | $\Delta S_m$ (kcal/K*mol) |
|---|---|---|---|
| AcpP | 57.5 ± 0.3 | 35.0 ± 2.0 | 0.60 ± 0.1 |
| ACT | 39.7 ± 1.7 | 26.6 ± 2.7 | 0.67 ± 0.1 |
| ACT$\mathcal{N}$-$\alpha_2$Ec | 30.9 ± 3.1 | 20.3 ± 5.0 | 0.64 ± 0.1 |
| ACT$\alpha_2$Ec | 53.7 ± 0.3 | 39.7 ± 5.5 | 0.74 ± 0.1 |
| Ec$\mathcal{N}$-$\alpha_2$ACT | - | - | - |
| Ec$\alpha_2$ACT | - | - | - |

**Table S5.** Quantitative analysis of AcpP-FabF and ACT-FabF RMSD simulated complexes.

| ACP | KS | Average Complex RMSD (Å) | Complex RSMD St. Dev. (Å) | Average Ppant RMSD (Å) | Ppant RMSD St. Dev. (Å) |
|---|---|---|---|---|---|
| AcpP | FabF | 3.446 | 0.443 | 3.210 | 0.575 |
| ACT | FabF | 4.366 | 0.619 | 7.356 | 3.302 |

**Table S6.** Amino acid sequence of AcpP with residues predicted to interact with FabF via salt bridges highlighted. Green = AcpP amino acids interacting in with FabF via salt bridges that do not have homologs in the ACT-FabF complex. Magenta = AcpP amino acids interacting with FabF via salt bridges that have homologs in the ACT-FabF complex.

Tens 1 2 3 4 5 6 7
Ones
123456789012345678901234567890123456789012345678901234567890123456789012345678
AcpP Sequence:
MSTIEERVKKIIGEQLGVKQEEVTNNASFVEDLGADSLDTVELVMALEEEFDTEIPDEEAEKITTVQAAIDYINGHQA

**Table S7.** Salt bridges in the AcpP-FabF simulated complex.

| Residue (ACP-KS) | Percentage (%) | Average Distance (Å) | St. Dev. (Å) |
|---|---|---|---|
| Glu48-Arg127 | 88.123 | 3.867 | 1.35 |
| Asp39-Lys65 | 79.502 | 3.645 | 1.481 |
| Glu49-Lys128 | 75.287 | 4.149 | 1.588 |
| Glu42-Lys69 | 56.705 | 4.704 | 2.359 |
| Asp57-Arg206 | 54.981 | 4.626 | 1.463 |
| Asp36-Lys65 | 43.103 | 4.960 | 1.763 |
| Glu42-Arg68 | 30.651 | 6.639 | 2.742 |
| Asp39-Lys69 | 15.900 | 5.980 | 1.702 |
| Asp36-Lys69 | 10.153 | 8.346 | 2.639 |
| Glu48-Lys128 | 8.812 | 6.918 | 1.907 |
| Asp36-His268 | 8.238 | 7.061 | 1.956 |
| Asp57-Arg127 | 5.939 | 5.985 | 1.355 |
| Glu54-Lys128 | 2.299 | 9.981 | 2.889 |
| Glu14-Arg68 | 1.916 | 8.938 | 3.005 |
| Glu14-Arg64 | 1.724 | 12.162 | 4.308 |
| Asp39-Arg68 | 1.149 | 8.632 | 2.382 |
| Glu54-Arg127 | 0.766 | 7.000 | 1.686 |
| Glu58-Arg211 | 0.476 | 14.918 | 3.207 |
| Asp71-His76 | 0.192 | 8.750 | 2.967 |
| Glu42-Lys65 | 0.192 | 9.015 | 2.003 |
| Glu61-Arg206 | 0.192 | 9.574 | 2.606 |

**Table S8.** Salt bridges in the ACT-FabF simulated complex.

| Residue (ACP-KS) | Percentage (%) | Average Distance (Å) | St. Dev. (Å) |
|---|---|---|---|
| Glu53-Lys128 | 68.966 | 4.214 | 2.439 |
| Glu47-Lys69 | 50.958 | 6.599 | 4.261 |
| Asp22-Lys65 | 41.954 | 6.581 | 4.019 |
| Asp63-Arg206 | 30.268 | 6.309 | 2.837 |
| Asp63-Arg127 | 13.602 | 7.074 | 2.454 |
| Asp41-Lys65 | 10.536 | 9.777 | 4.562 |
| Glu53-Arg127 | 8.621 | 10.418 | 3.368 |
| Asp41-Lys69 | 8.621 | 10.285 | 4.047 |
| Asp22-Arg64 | 7.663 | 12.650 | 5.897 |
| Asp41-Arg206 | 6.322 | 13.108 | 3.193 |
| Asp41-His268 | 3.831 | 8.289 | 2.397 |
| Glu20-Lys65 | 3.448 | 10.694 | 3.327 |
| Asp41-Arg68 | 2.299 | 13.100 | 4.622 |
| Glu47-Arg68 | 1.149 | 10.600 | 3.119 |
| Glu20-Arg64 | 0.958 | 10.982 | 4.246 |
| Arg51-Glu116 | 0.958 | 9.099 | 2.628 |
| Arg55-Glu116 | 0.766 | 12.759 | 3.48 |
| Asp62-Arg127 | 0.575 | 9.655 | 3.154 |
| Glu16-Arg68 | 0.575 | 10.518 | 3.631 |
| Glu16-Arg64 | 0.383 | 13.964 | 4.438 |
| Glu47-Lys128 | 0.192 | 11.857 | 1.658 |
| Asp62-Arg206 | 0.192 | 12.784 | 2.549 |

**Table S9.** Salt bridges in the ACT$\alpha_2$Ec-FabF simulated complex.

| Residue (ACP-KS) | Percentage (%) | Average Distance (Å) | St. Dev. (Å) |
|---|---|---|---|
| Glu47-Lys69 | 99.234 | 3.172 | 0.485 |
| Asp63-Lys128 | 72.797 | 4.485 | 2.976 |
| Glu47-Lys65 | 51.341 | 5.319 | 2.222 |
| Glu53-Arg68 | 50.000 | 4.900 | 2.099 |
| Asp62-Lys128 | 45.211 | 5.717 | 2.876 |
| Glu54-Arg64 | 15.326 | 7.142 | 2.815 |
| Glu54-Lys65 | 12.452 | 9.762 | 3.413 |
| Glu53-Arg64 | 12.452 | 9.870 | 4.106 |
| Glu20-Lys65 | 11.877 | 12.480 | 6.395 |
| Asp44-His268 | 11.877 | 5.350 | 1.110 |
| Asp69-Arg206 | 3.640 | 8.954 | 2.569 |
| Glu54-Arg68 | 1.533 | 7.803 | 2.513 |
| Asp63-Arg127 | 1.533 | 9.427 | 2.328 |
| Glu36-Arg206 | 0.958 | 9.386 | 2.192 |
| Glu54-Lys128 | 0.575 | 19.178 | 2.948 |
| Glu55-Lys65 | 0.383 | 12.372 | 2.426 |
| Glu47-Arg68 | 0.383 | 10.465 | 1.728 |
| Glu20-Arg64 | 0.383 | 20.177 | 6.940 |
| Asp62-Arg68 | 0.192 | 8.839 | 2.972 |
| Asp62-Arg206 | 0.192 | 12.677 | 2.241 |

**Table S10.** List of theoretical and experimental MW of the chimeric ACPs. Theoretical MWs were calculated based on the amino acid sequence of each chimera. Experimental MWs were obtained by deconvoluting the LCMS results (Electrospray mass spectra, **Figure S1–S9**) using ESIprot.

| Fig # | Protein | Theoretical MW of pre-modified protein (Da) | Theoretical MW of post-modified protein (Da) | Experimental (deconvoluted) MW (Da) | Correct product ? |
|---|---|---|---|---|---|
| S1A | *holo*-AcpP | *apo*: 10802.8 | 11010.3 | 11010.8 ± 0.4 | yes |
| S1B | *holo*-AcpP-TNB | *holo*: 11010.8 | 11207.9 | 11209.1 ± 0.5 | yes |
| S2A | *holo*-ACT | *apo*: 11395.5 | 11603.6 | 11603.6 ± 0.3 | yes |
| S2B | *holo*-ACT-TNB | *holo*: 11603.5 | 11800.6 | 11800.7 ± 0.3 | yes |
| S3A | *holo*-ACT*N*-$\alpha_1$Ec | *apo*: 11293.5 | 11501.6 | 11501.6 ± 0.4 | yes |
| S3B | *holo*-ACT*N*-$\alpha_1$Ec-TNB | *holo*: 11501.6 | 11698.6 | 11698.2 ± 0.5 | yes |
| S4A | *holo*-ACT*N*-$\alpha_2$Ec | *apo*: 10937.1 | 11145.2 | 11144.8 ± 0.5 | yes |
| S4B | *holo*-ACT*N*-$\alpha_2$Ec-TNB | *holo*: 11144.8 | 11342.2 | 11342.2 ± 0.4 | yes |
| S5A | *holo*-ACT$\alpha_2$Ec | *apo*: 11409.5 | 11617.6 | 11617.6 ± 0.5 | yes |
| S5B | *holo*-ACT$\alpha_2$Ec-TNB | *holo*: 11617.7 | 11814.6 | 11814.7 ± 0.7 | yes |
| S6A | *holo*-ACT$l_2$Ec | *apo*: 11497.6 | 11705.7 | 11705.6 ± 0.6 | yes |
| S6B | *holo*-ACT$l_2$Ec-TNB | *holo*: 11705.6 | 11902.7 | 11902.5 ± 0.7 | yes |
| S7A | *holo*-ACT$l_2$-$l_3$Ec | *apo*:11585.7 | 11793.8 | 11793.5 ± 0.6 | yes |
| S7B | *holo*-ACT$l_2$-$l_3$Ec-TNB | *holo*: 11793.5 | 11990.8 | 11990.0 ±0.6 | yes |
| S8A | *holo*-Ec*N*-$\alpha_2$ACT | *apo*: 11261.4 | 11469.5 | 11469.3 ± 0.5 | yes |
| S8B | *holo*-Ec*N*-$\alpha_2$ACT-TNB | *holo*: 11469.3 | 11666.5 | 11666.3 ± 0.4 | yes |
| S9A | *holo*-Ec$\alpha_2$ACT | *apo*: 10788.9 | 10997.0 | 10996.6 ± 0.3 | yes |
| S9B | *holo*-Ec$\alpha_2$ACT-TNB | *holo*: 10996.6 | 11194.0 | 11194.1 ± 0.3 | yes |

**Figure S1–S9.** LCMS results: Electrospray (ES) mass spectra for the ACPs used in this study (AcpP, ACT, the AcpP/ACT chimeras, post-Sfp reaction and post-DTNB reaction). For each ES mass spectrum, the theoretical MW is indicated along with the MWs of the deconvoluted peaks. The change in MW (ΔMW) and the modifications for each species are indicated. Some proteins lost their *N*-terminal methionine and some exhibited minor gluconoylation, both of which are not expected to interfere with other analyses.

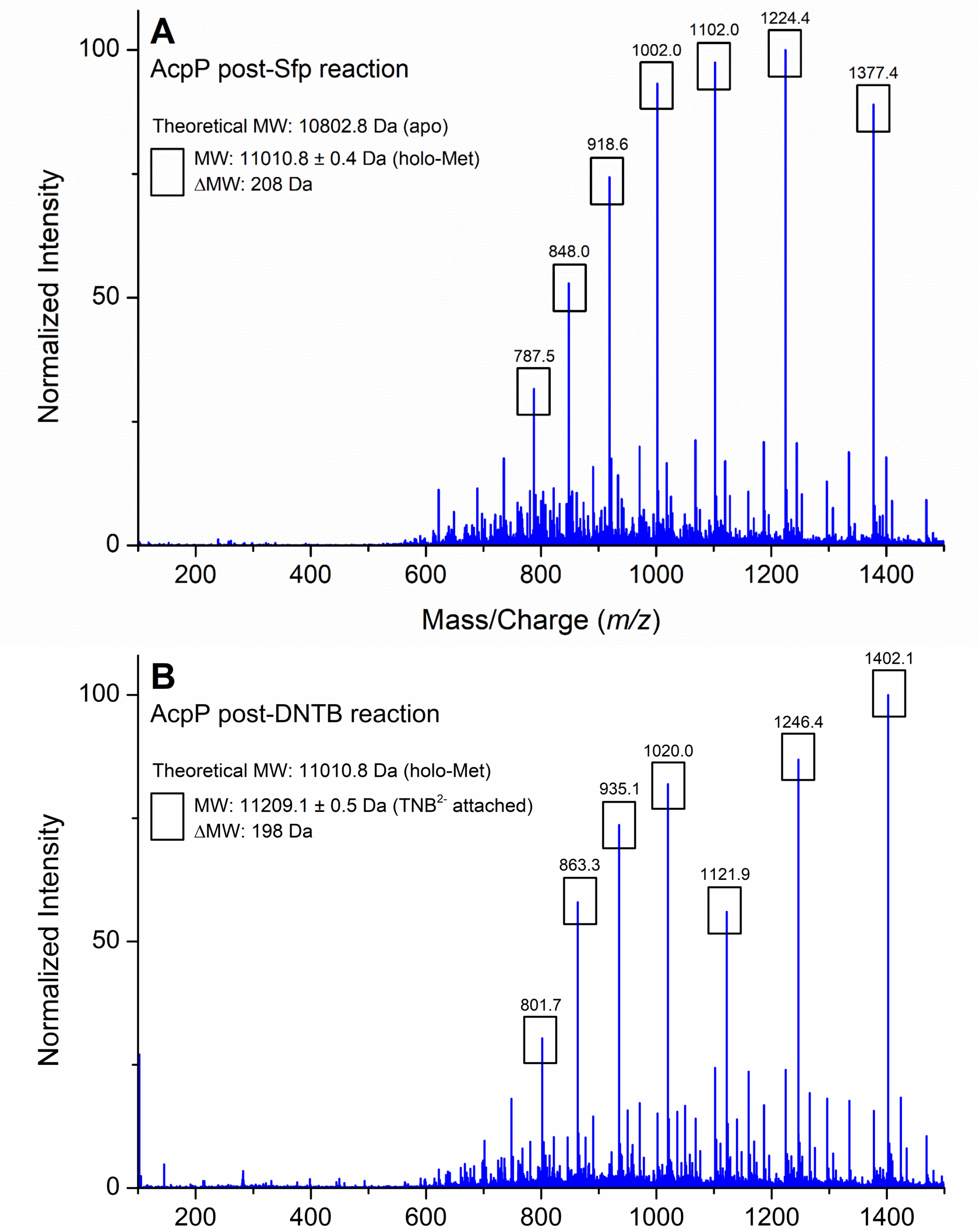


**Figure S1.** ES mass spectrum of post-Sfp reaction *holo* AcpP in (A) and post-DTNB reaction AcpP-TNB in (B). The theoretical MW is compared to the deconvoluted MW and the ΔMW is noted. Results indicate a 100% conversion of *apo* to *holo* and successful attachment of the $TNB^{2-}$ probe.

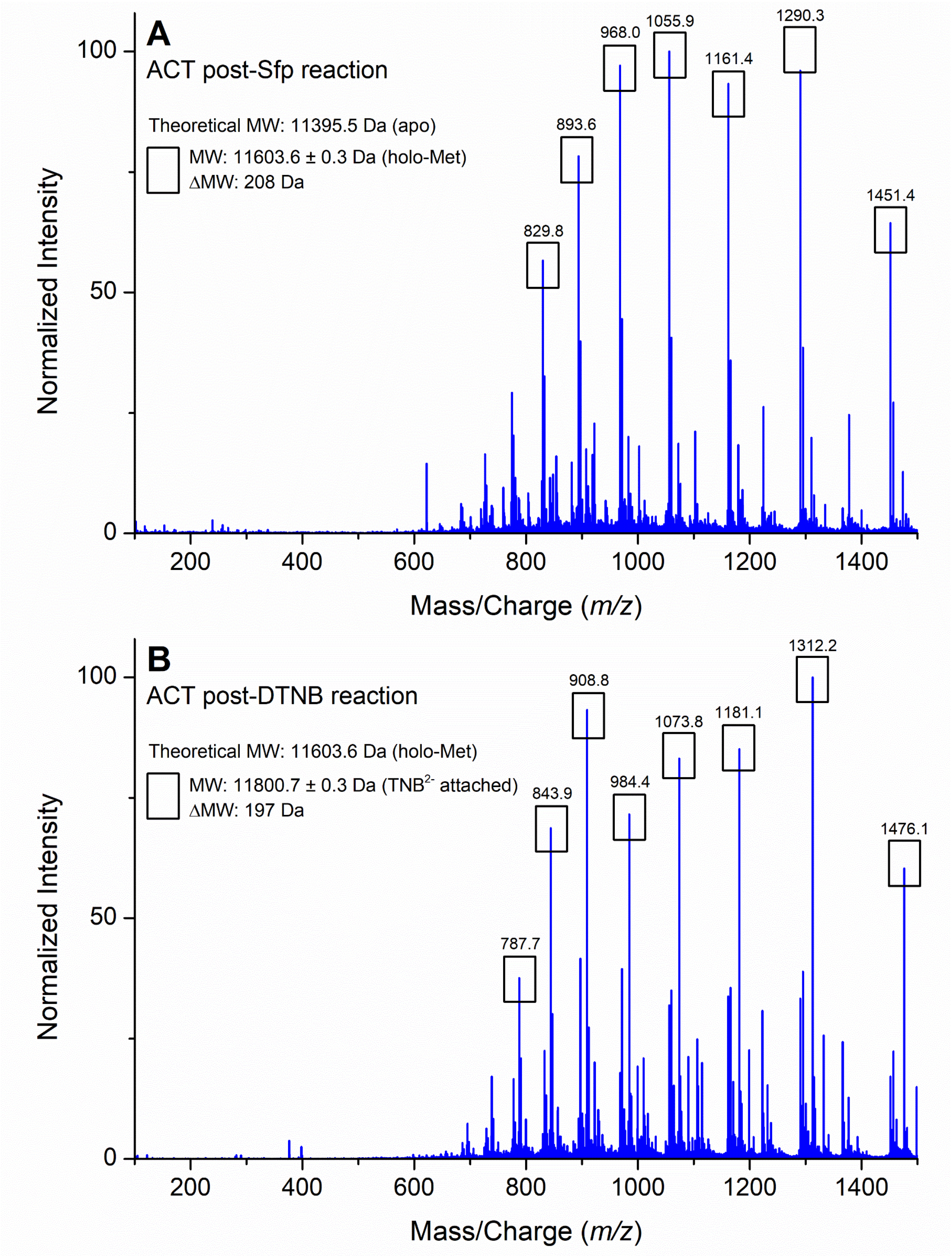


**Figure S2.** ES mass spectrum of post-Sfp reaction *holo* ACT in (A) and post-DTNB reaction ACT-TNB in (B). The theoretical MW is compared to the deconvoluted MW and the ΔMW is noted. Results indicate a 100% conversion of *apo* to *holo* and successful attachment of the $TNB^{2-}$ probe.

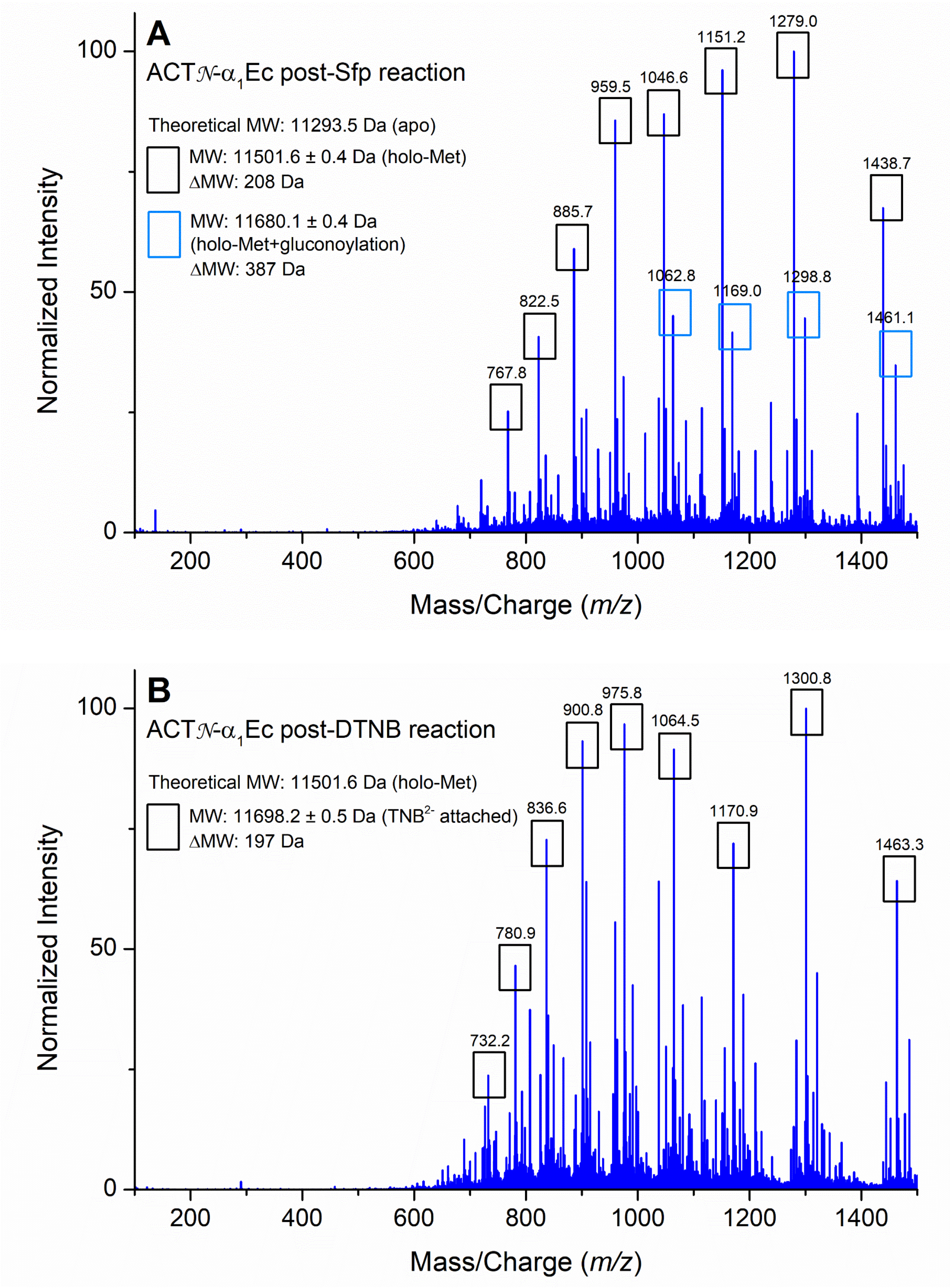


**Figure S3.** ES mass spectrum of post-Sfp reaction *holo* ACT𝒩-$\alpha_1$Ec in (A) and post-DTNB reaction ACT𝒩-$\alpha_1$Ec-TNB in (B). The theoretical MW is compared to the deconvoluted MW and the ΔMW is noted. Results indicate a 100% conversion of *apo* to *holo* and successful attachment of the $TNB^{2-}$ probe.

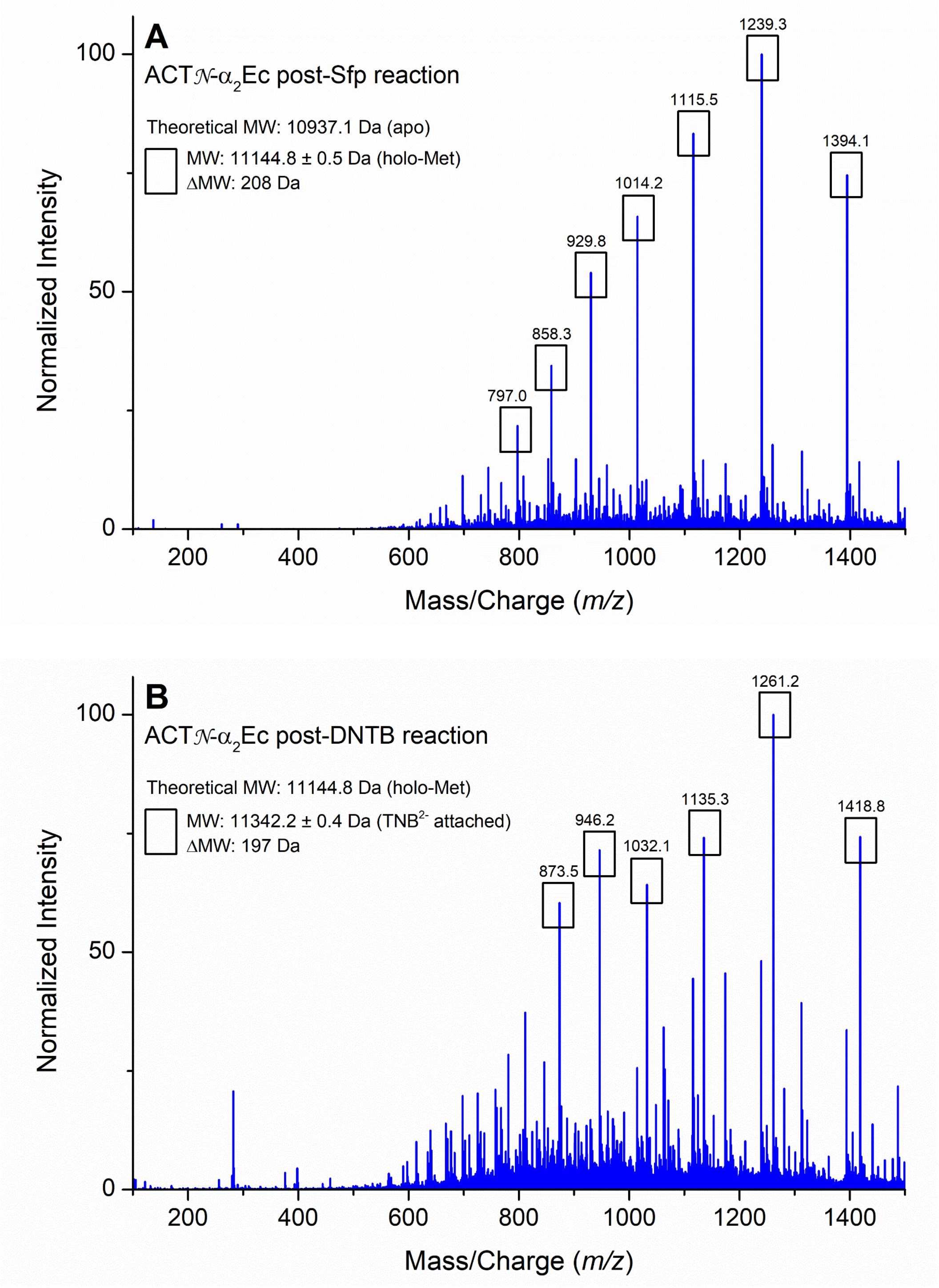


**Figure S4.** ES mass spectrum of post-Sfp reaction *holo* ACT𝒩-$\alpha_2$Ec in (A) and post-DTNB reaction ACT𝒩-$\alpha_2$Ec-TNB in (B). The theoretical MW is compared to the deconvoluted MW and the ΔMW is noted. Results indicate a 100% conversion of *apo* to *holo* and successful attachment of the $TNB^{2-}$ probe.

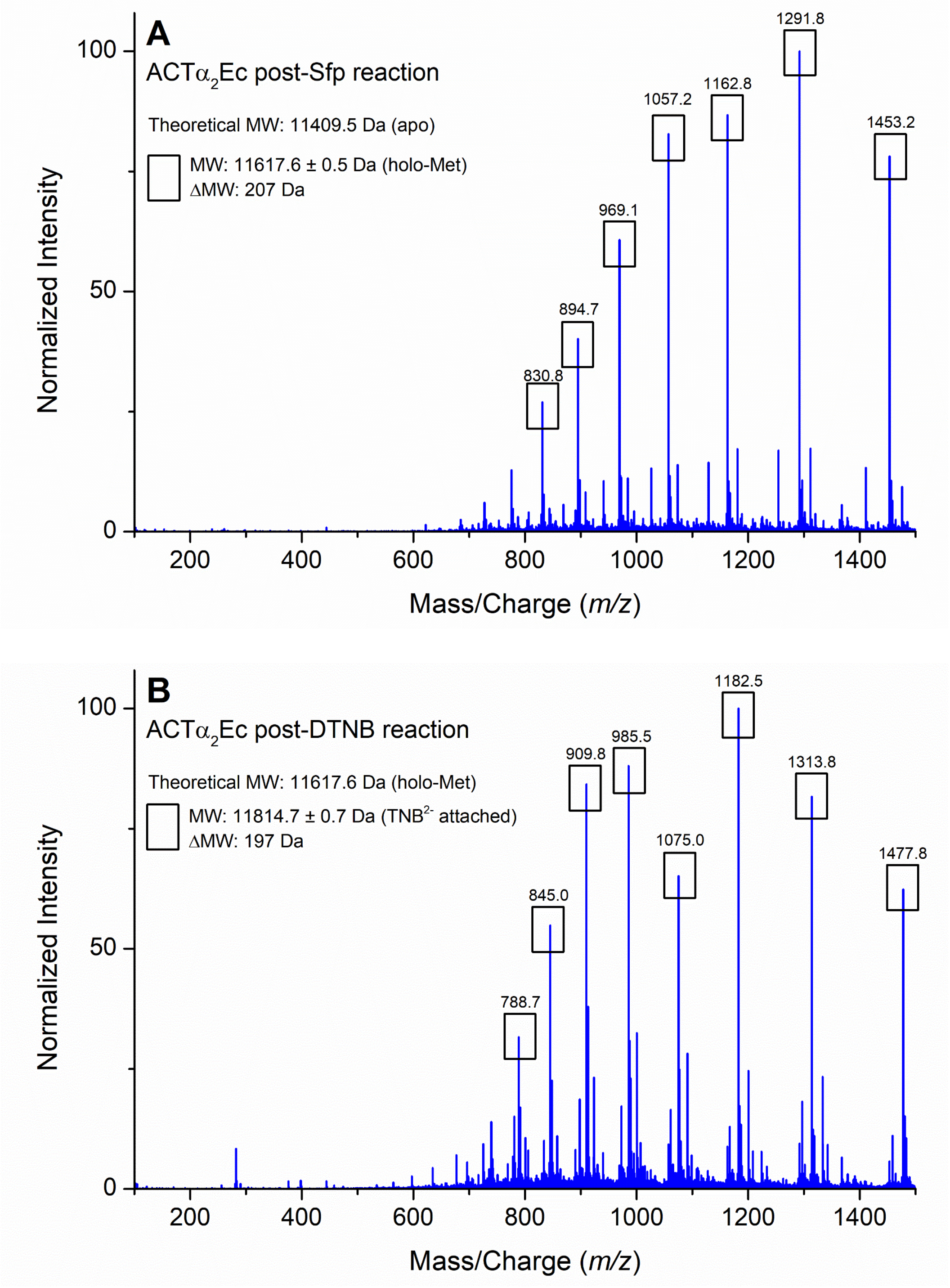


**Figure S5.** ES mass spectrum of post-Sfp reaction *holo* ACTα2Ec in (A) and post-DTNB reaction ACTα2Ec-TNB in (B). The theoretical MW is compared to the deconvoluted MW and the ΔMW is noted. Results indicate a 100% conversion of *apo* to *holo* and successful attachment of the $TNB^{2-}$ probe.

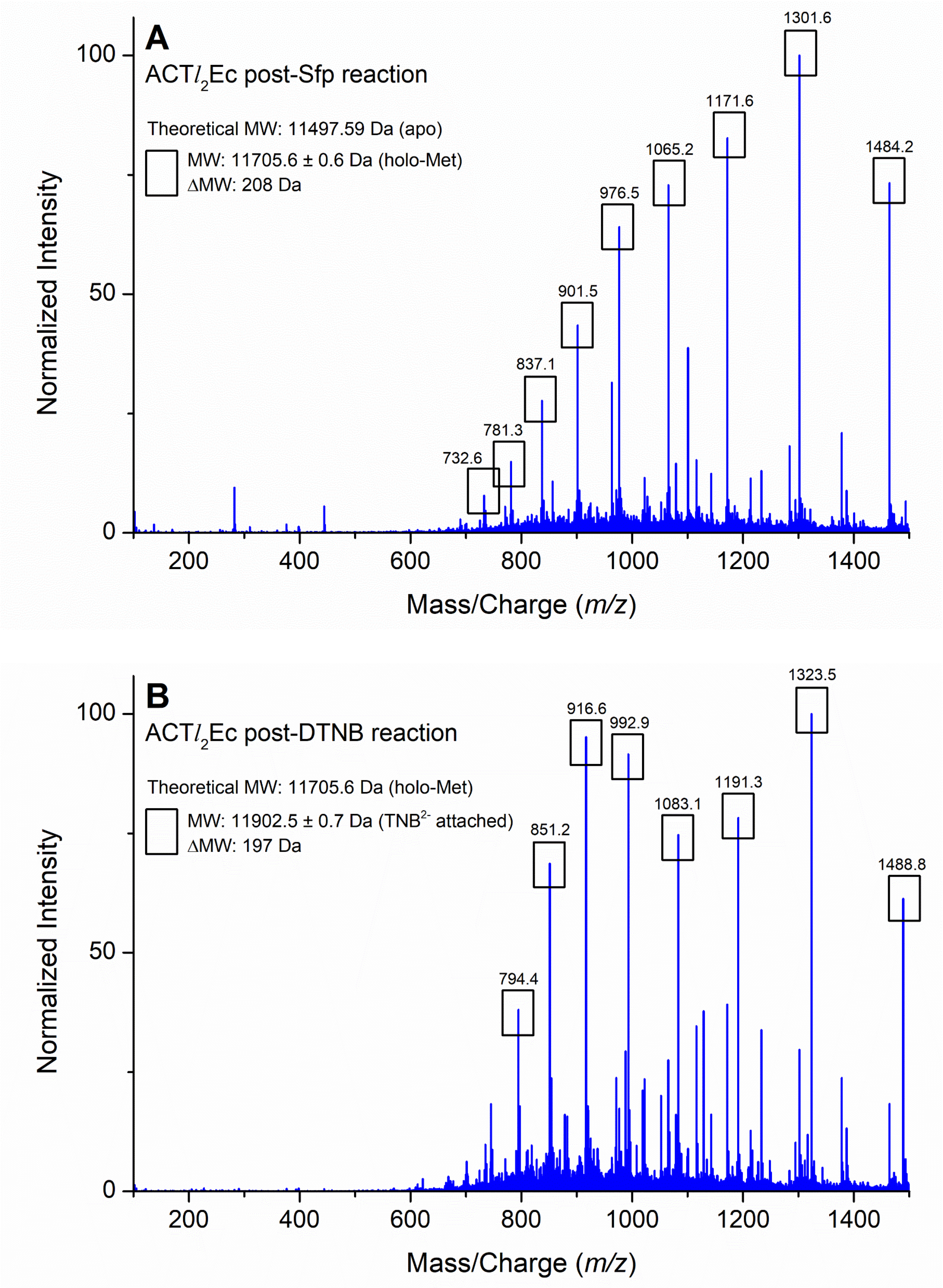


**Figure S6.** ES mass spectrum of post-Sfp reaction *holo* ACT*l*2Ec in (A) and post-DTNB reaction ACT*l*2Ec-TNB in (B). The theoretical MW is compared to the deconvoluted MW and the ΔMW is noted. Results indicate a 100% conversion of *apo* to *holo* and successful attachment of the $TNB^{2-}$ probe.

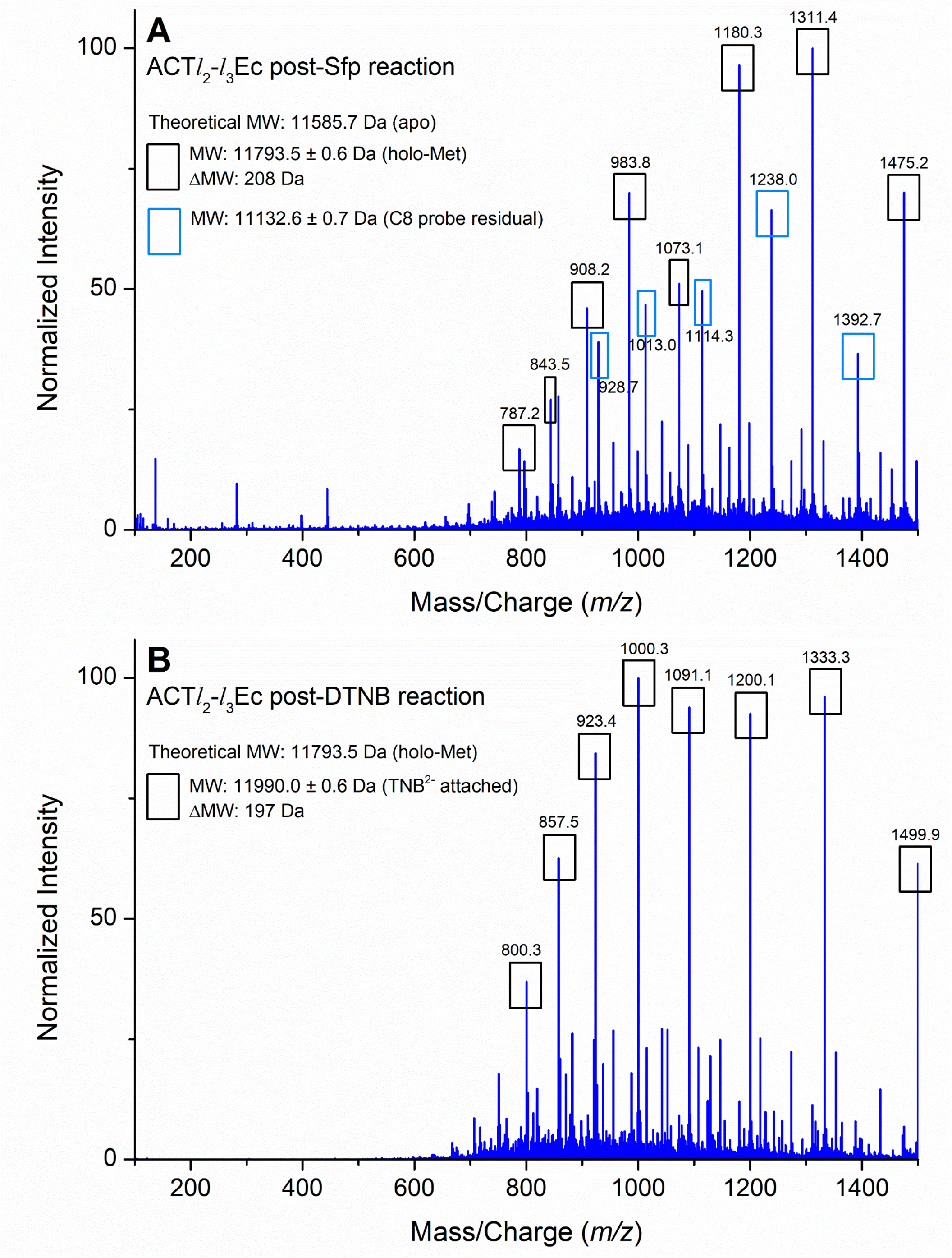


**Figure S7.** ES mass spectrum of post-Sfp reaction *holo* ACT$l_2$-$l_3$Ec in (A) and post-DTNB reaction ACT$l_2$-$l_3$Ec-TNB in (B). The theoretical MW is compared to the deconvoluted MW and the ΔMW is noted. Results indicate a 100% conversion of *apo* to *holo* and successful attachment of the $TNB^{2-}$ probe.

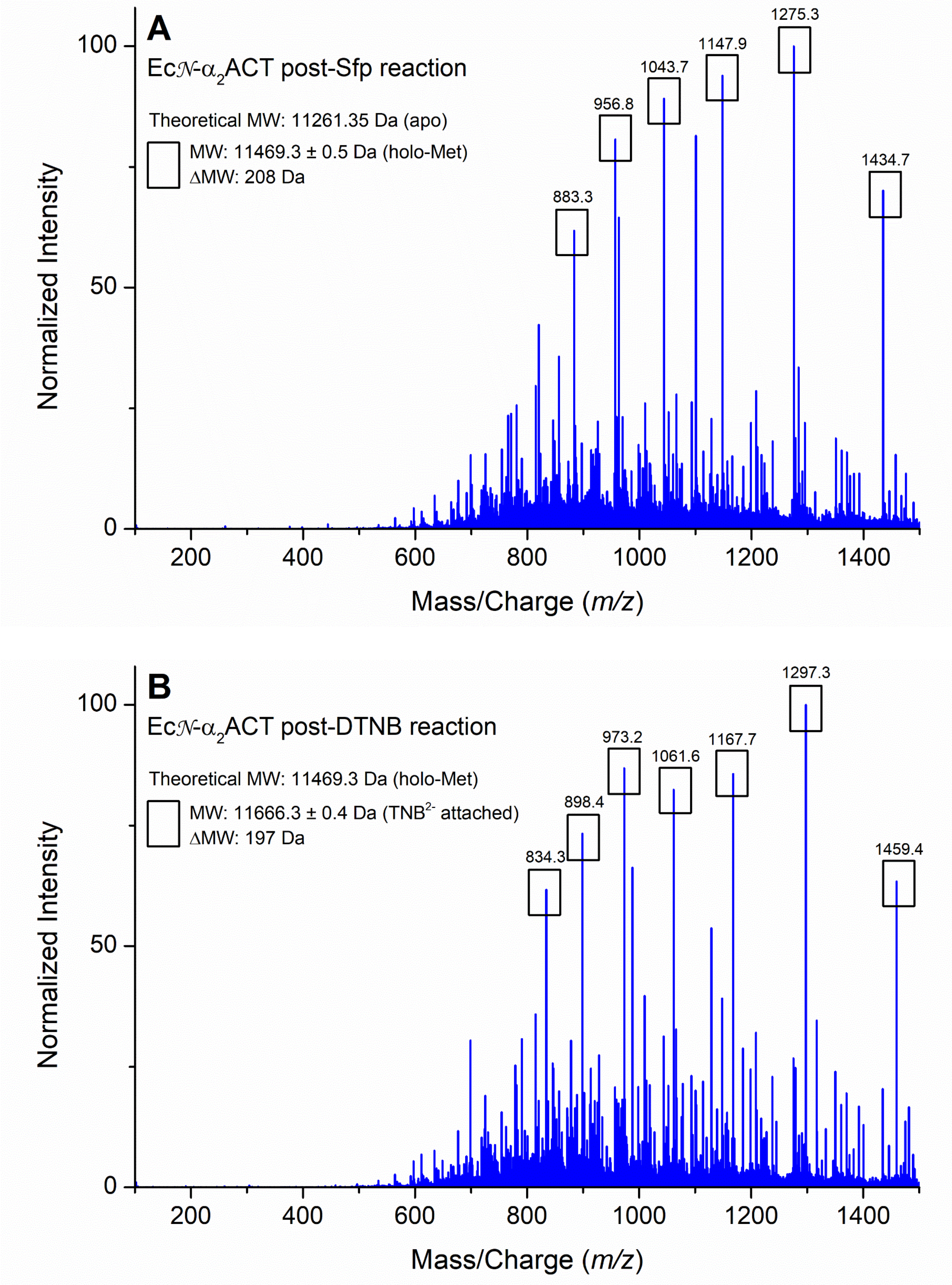


**Figure S8.** ES mass spectrum of post-Sfp reaction *holo* Ec$\mathcal{N}$-$\alpha_2$ACT in (A) and post-DTNB reaction Ec$\mathcal{N}$-$\alpha_2$ACT-TNB in (B). The theoretical MW is compared to the deconvoluted MW and the ΔMW is noted. Results indicate a 100% conversion of *apo* to *holo* and successful attachment of the $TNB^{2-}$ probe.

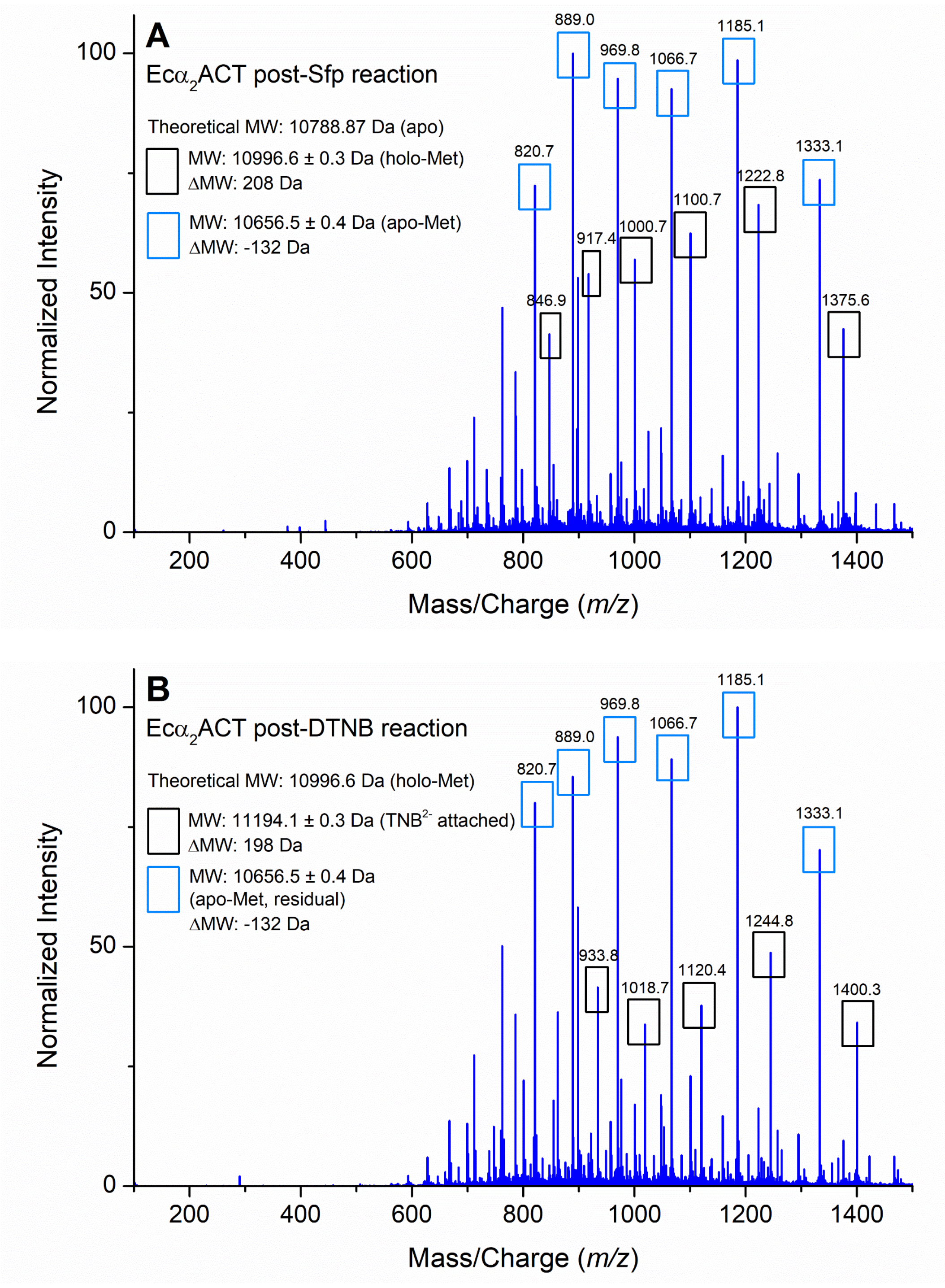


**Figure S9.** ES mass spectrum of post-Sfp reaction *holo* Ecα$_2$ACT in (A) and post-DTNB reaction Ecα$_2$ACT-TNB in (B). The theoretical MW is compared to the deconvoluted MW and the ΔMW is noted. Results indicate a 40% conversion of *apo* to *holo* and successful attachment of the TNB$^{2-}$ probe.

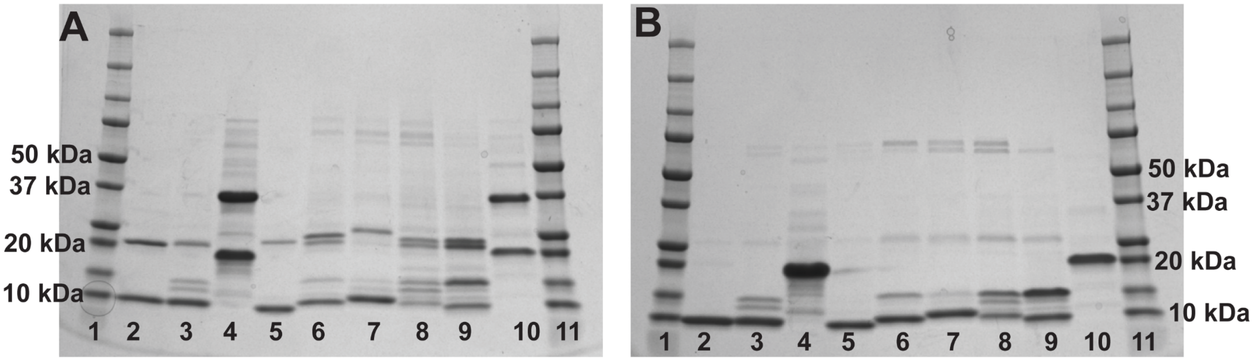


**Figure S10.** Analysis of ACPs in this study by SDS PAGE under non-reducing (A) and reducing (B) conditions. Most ACP chimeras exhibit migration patterns that are similar to that of their template ACT, however ACT$\mathcal{N}$-$\alpha_2$Ec shows AcpP-like migration. The addition of + 5% (v/v) BME under reducing conditions (B) breaks apart ACP dimers that are observed under non-reducing conditions (A). Lane 1. Protein ladder; Lane 2. ACT; Lane 3. ACT$\mathcal{N}$-$\alpha_1$Ec; Lane 4. ACT$\mathcal{N}$-$\alpha_2$Ec; Lane 5. ACT$\alpha_2$Ec; Lane 6. ACT$l_2$Ec; Lane 7. ACT$l_2$-$l_3$Ec; Lane 8. ACT$\alpha_3$-$\alpha_4$Ec; Lane 9. ACT$\alpha_4$Ec; Lane 10. AcpP; Lane 11. Protein ladder.

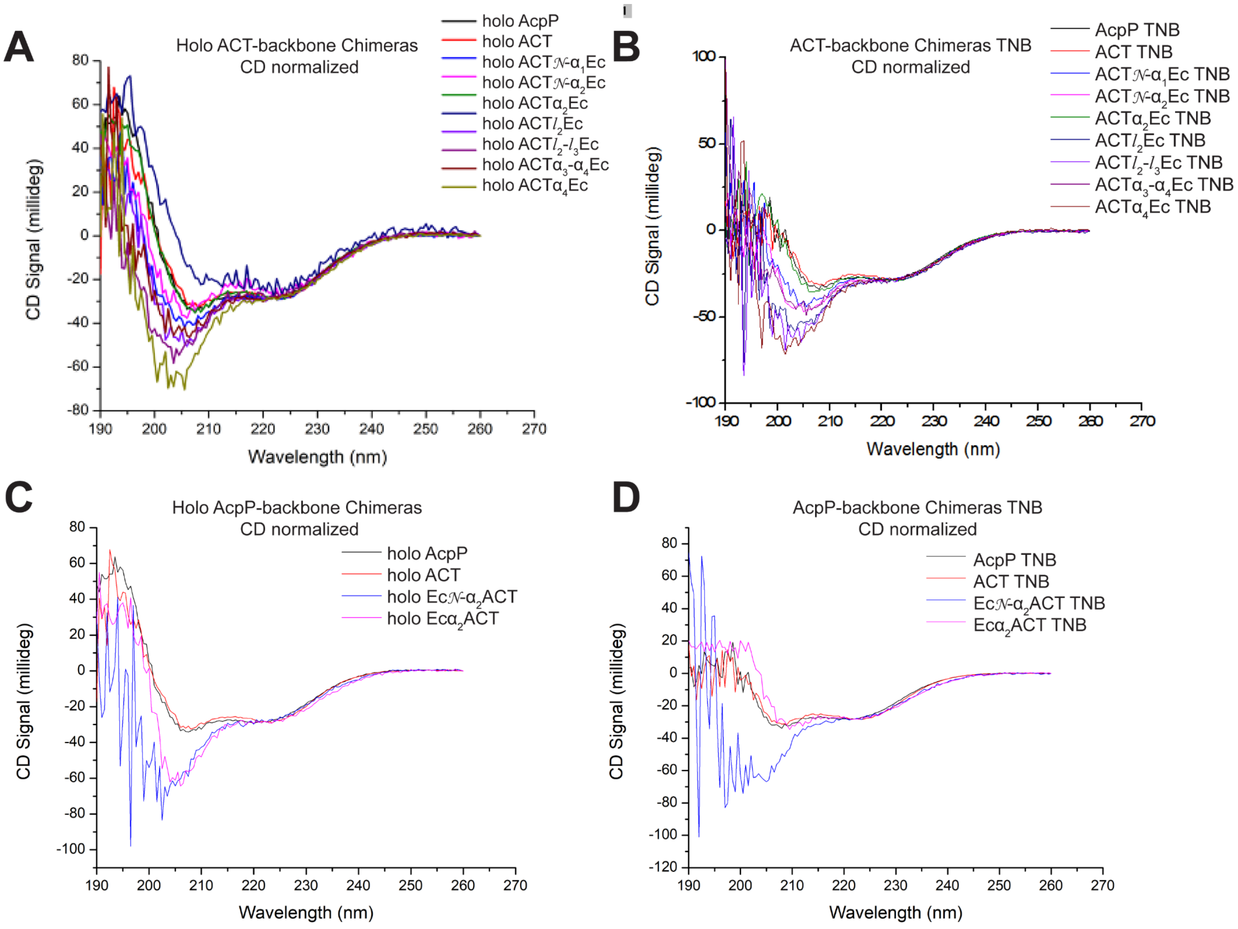


**Figure S11.** CD Spectra of *holo* (A-B) and post-DTNB reaction TNB-attached (C-D) ACPs used in this study. (A) Comparison of the ACT-backbone chimeras to AcpP and ACT, all in their *holo* forms. (B) Comparison of the ACT-backbone chimeras to AcpP and ACT, all in their TNB-attached forms. (C) Comparison of the AcpP-backbone chimeras to AcpP and ACT, all in their *holo* forms. (D) Comparison of the AcpP-backbone chimeras to AcpP and ACT, all in their TNB-attached forms.

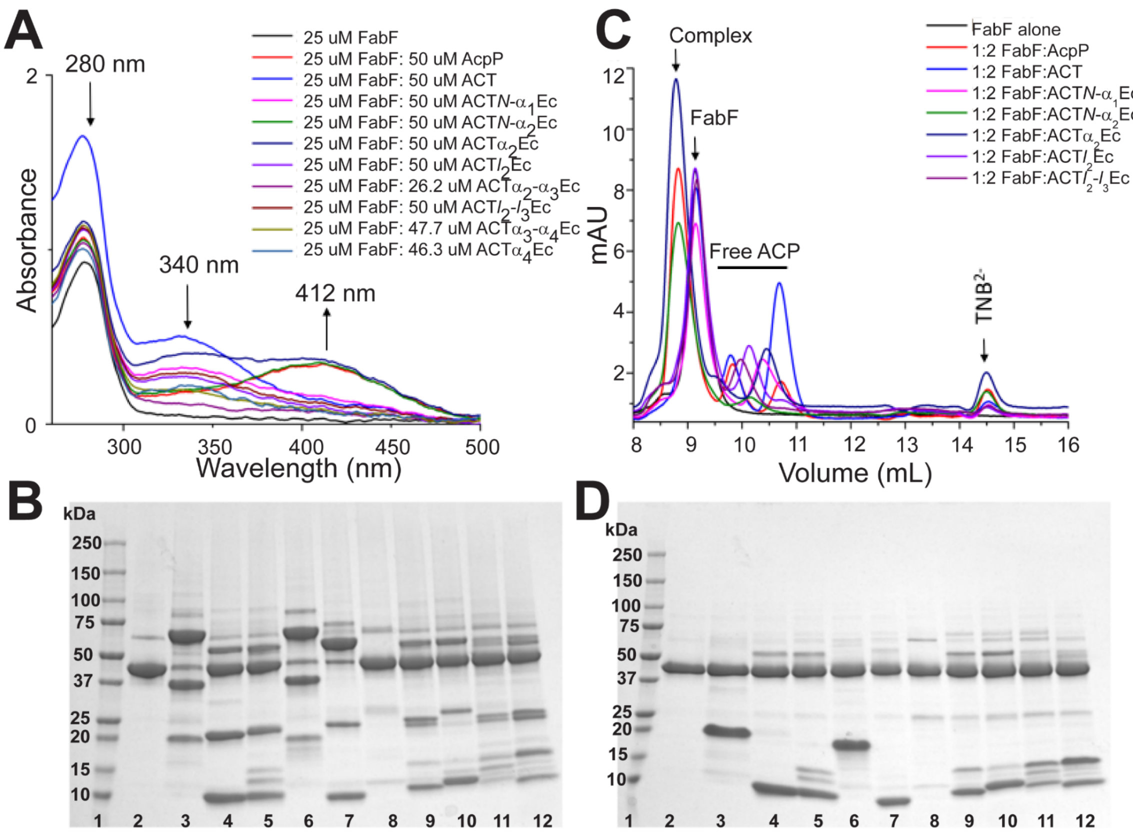


**Figure S12.** Analysis of ACT-backbone chimeric ACP binding to FabF via a colorimetric mechanism based crosslinking assay. (A) UV-Vis spectra of ACT-backbone chimera crosslinking assays**.** The peak at 280 nm corresponds to *holo* ACP, the peak at 340 nm corresponds to ACP-$TNB^{-}$, and the peak at 412 nm corresponds to the released $TNB^{2-}$ ion. The presence of a 412 nm peak in the AcpP, ACT$\mathcal{N}$-$\alpha_2$Ec, and ACT$\alpha_2$Ec samples indicates that these ACPs successfully crosslinked to FabF causing the release of $TNB^{2-}$. (C) Size exclusion chromatography (SEC) chromatogram of ACT-backbone chimera crosslinking assays. The ACP-FabF complex elutes first at 9 mL followed by FabF which elutes at 10–11 mL. The standalone ACPs elute between 12–14 mL, and lastly $TNB^{2-}$ elutes at 21 mL. The ACPs that crosslinked successfully to FabF (AcpP, ACT$l_1$-$\alpha_2$Ec, and ACT$\alpha_2$Ec) can be observed by the presence of a higher molecular weight complex and $TNB^{2-}$, as well as the absence of standalone FabF. (A & D) SDS PAGE gel of ACT-backbone chimera crosslinking assays. (B) Non-reducing gel of ACP-FabF crosslinking assay. ACP-$TNB^{-}$ monomer and dimers at ~10–40 kDa, FabF at ~45 kDa, and ACP-FabF complex at ~70 kDa. FabF depletion and a higher molecular weight band is present in Lanes 3, 6, and 7, indicating that ACP-FabF crosslinking occurred.

Lane 1. Protein ladder; Lane 2. FabF only; Lane 3. AcpP + FabF; Lane 4. ACT + FabF; Lane 5. ACT$l_1$-$\alpha_1$Ec + FabF; Lane 6. ACT$l_1$-$\alpha_2$Ec + FabF; Lane 7. ACT$\alpha_2$Ec + FabF; Lane 8. ACT$\alpha_2$-$\alpha_3$Ec + FabF; Lane 9. ACT$l_3$Ec + FabF; Lane 10. ACT$l_3$-$l_4$Ec + FabF; Lane 11. ACT$\alpha_3$-$\alpha_4$Ec + FabF; Lane 12. ACT$\alpha_4$Ec + FabF. (D) Reducing gel of ACP-FabF crosslinking assay. All crosslinks are reduced upon the addition of BME resulting in the loss of higher molecular weight complex bands ~70 kDa and recovery of FabF concentration ~45 kDa. All ACP-$TNB^{-}$ dimers were reduced to monomers. Same Lanes 1–12 as detailed in (B).

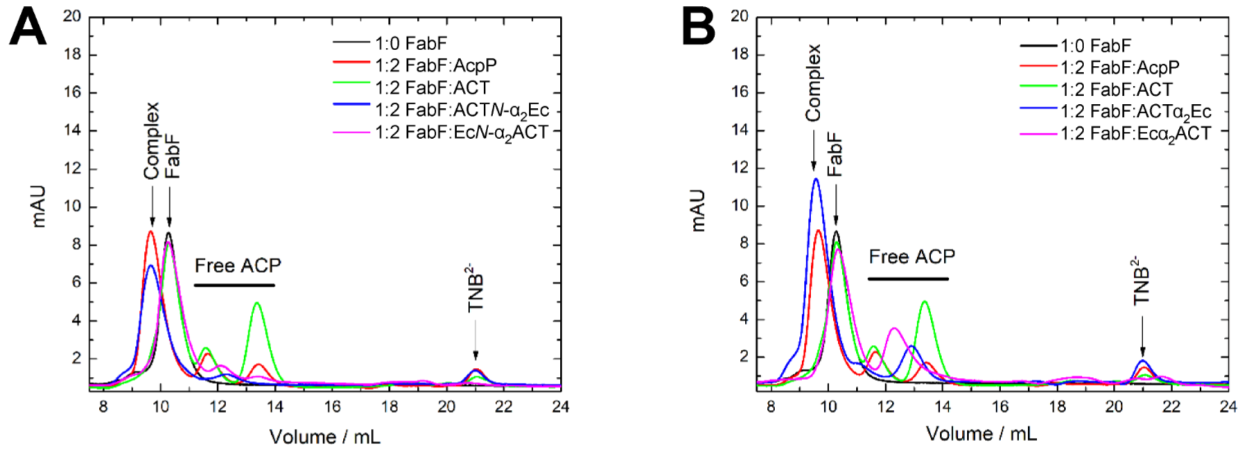


**Figure S13.** Size exclusion chromatography results showing Absorbance at 280 nm. When AcpP-TNB⁻, ACT$\mathcal{N}$-α$_2$Ec-TNB⁻ (A), and ACTα$_2$Ec-TNB⁻ (B) are mixed with FabF, a higher molecular weight complex is formed (9.5 minutes) and a smaller molecule TNB$^{2-}$ (21 min) is released. Conversely, when ACT, Ec$\mathcal{N}$-α$_2$ACT-TNB⁻ and Ec$\mathcal{N}$-α$_2$ACT-TNB⁻ are mixed with FabF, neither the band corresponding to the higher molecular weight complex nor the release of TNB$^{2-}$ are observed. Instead, bands corresponding to stand alone FabF (10 min) and the ACP (12–14 minutes) are observed.

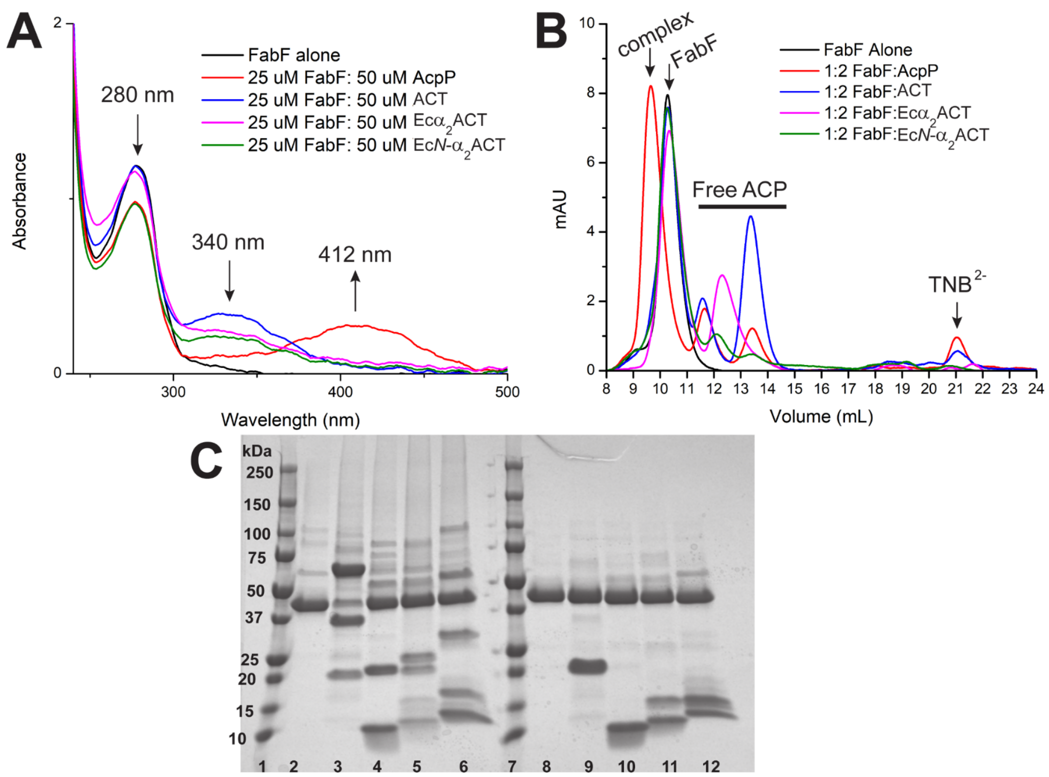


**Figure S14.** Analysis of AcpP-backbone chimeric ACP binding to FabF via a colorimetric mechanism based crosslinking assay. (A) UV-Vis analysis of mechanism based crosslinking reaction mixtures. The peak at 280 nm corresponds to *holo* ACP, the peak at 340 nm corresponds to ACP-$TNB^{-}$ attached, and the peak at 412 nm corresponds to the $TNB^{2-}$ ion. The presence of a 412 nm peak in the AcpP sample indicates that this ACP successfully crosslinked to FabF, causing the release of $TNB^{2-}$. The absence of a 412 nm peak in the Ec$\mathcal{N}$-$\alpha_2$ACT and Ec$\alpha_2$ACT samples indicates that chimeras did not crosslink to FabF. (B) Size exclusion chromatography (SEC) chromatogram of mechanism based crosslinking reaction mixtures. The ACP-FabF complex elutes first at 9 mL followed by FabF which elutes at 10–11 mL. The free ACPs elute anywhere between 12–14 mL depending on their molecular weight, and lastly $TNB^{2-}$ elutes at 21 mL. AcpP crosslinked successfully to FabF, resulting in a higher molecular weight complex that eluted first. In addition, this sample did not contain any FabF alone as it was all in complex with ACP, and had a peak for $TNB^{2-}$ which eluted at last. Ec$\mathcal{N}$-$\alpha_2$ACT and Ec$\alpha_2$ACT did not crosslink to FabF, as indicate by a lack of peaks corresponding to the ACP-FabF complex (9 mL) and $TNB^{2-}$ (21 mL) and presence of FabF alone (10–12 mL) and standalone ACP (12–14 mL). (C) SDS PAGE analysis of mechanism based crosslinking reaction mixtures under reducing and non-reducing conditions. The AcpP-$TNB^{2-}$ + FabF (lane 3) was the only mixture that generated a higher molecular weight complex ~70 kDa, indicating that the AcpP-backbone ACP chimeras studied lost their ability to interact with FabF. As expected, the ACT-$TNB^{-}$ did not form a complex with FabF. Upon addition of BME, the AcpP-FabF crosslink was reduced (lane 9), the FabF band ~45 kDa matched the concentration of the other mixtures. All ACP complexes were also reduced. Non-reducing lanes 2–6. Lane 1. Protein ladder; Lane 2. FabF only; Lane 3. AcpP-$TNB^{-}$ + FabF; Lane 4. Ec$\mathcal{N}$-$\alpha_2$ACT-$TNB^{-}$ + FabF; Lane 5. Ec$\alpha_2$ACT-$TNB^{-}$ + FabF; Lane 6. ACT-$TNB^{-}$ + FabF. Reducing lanes 8–12. Lane 7. Protein ladder; Lane 8. FabF only; Lane 9. AcpP-$TNB^{-}$ + FabF; Lane 10. Ec$\mathcal{N}$-$\alpha_2$ACT -$TNB^{-}$ + FabF; Lane 11. Ec$\alpha_2$ACT-$TNB^{-}$ + FabF; Lane 12. ACT-$TNB^{-}$ + FabF.

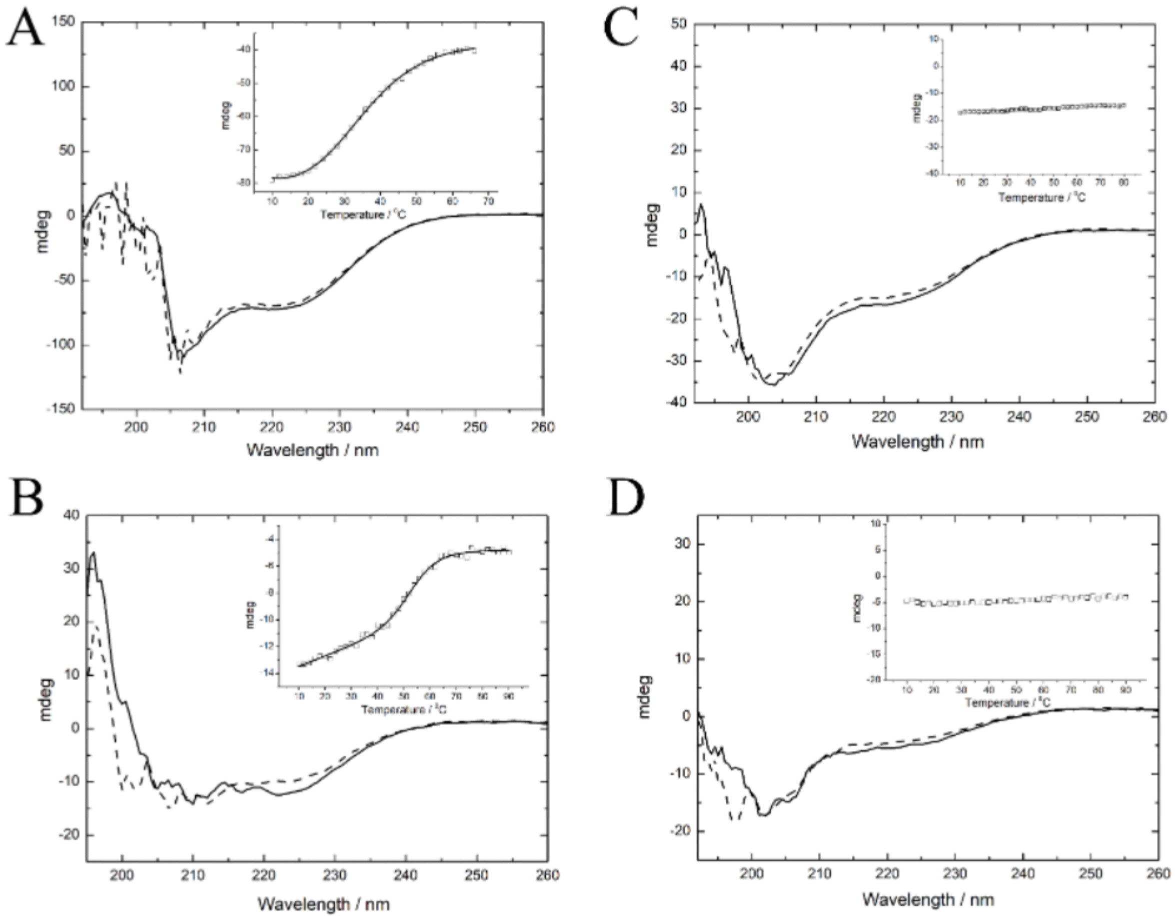


**Figure S15.** Thermal unfolding of ACP chimeras: ACT$\mathcal{N}$-$\alpha_2$Ec (A), ACT$\alpha_2$Ec (B), Ec$\mathcal{N}$-$\alpha_2$ACT (C), and Ec$\alpha_2$ACT (D). Full CD spectra of ACPs are shown for pre-$T_{melt}$ (solid lines) and post-$T_{melt}$ (dashed lines) samples. The inset shows $T_{melt}$ curves, with $T_m$ values obtained upon fitting the fractional change of unfolded chimeric ACP as a function of temperature to a N↔D model (N = native, D = denatured) with $\Delta Cp = 0$. Best fit data are presented as solid lines. See Table S4 for change in entropy and enthalpy of unfolding. ACT$\mathcal{N}$-$\alpha_2$Ec (A) and ACT$\alpha_2$Ec (B) show reversibility of unfolding Ec$\mathcal{N}$-$\alpha_2$ACT (C), and Ec$\alpha_2$ACT (D) show a general loss of secondary structure and no change in absorbance at 222 nm upon increase in temperature.

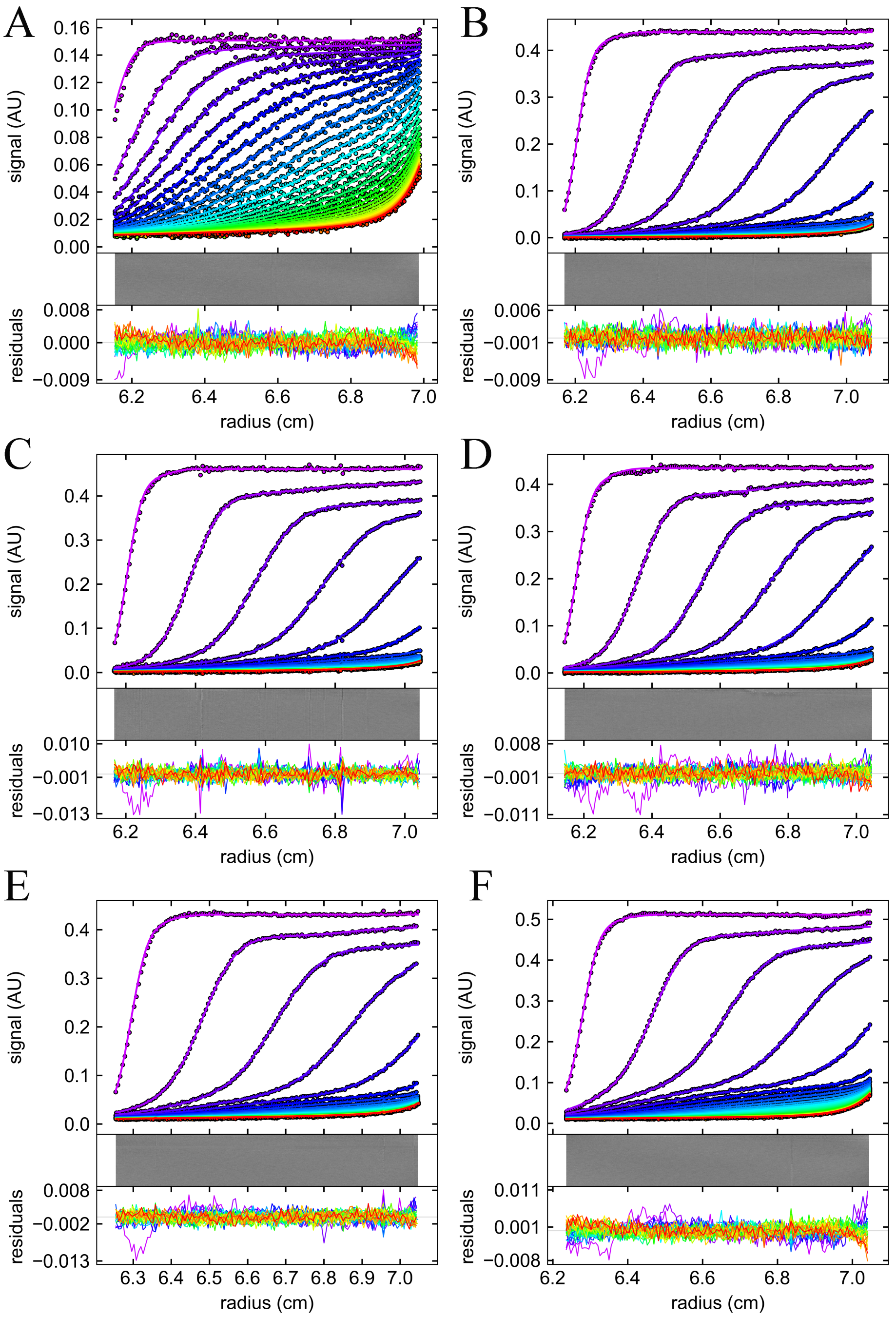


**Figure S16.** SV-AUC raw data and fits related to Figure 5. Representative subset, every 8-th scan, of the raw data and their fits after elimination of time invariant noise contributions and their residuals. The quality of fits to a c(s) model resulted in few diagonal features. (A) 150 μM ACT$\mathcal{N}$-$\alpha_2$Ec; (B) 17 μM of FabF with no ACT$\mathcal{N}$-$\alpha_2$Ec; (C) 17 μM FabF with 2.7 μM ACT$\mathcal{N}$-$\alpha_2$Ec; (D) 17 μM FabF with 5 μM ACT$\mathcal{N}$-$\alpha_2$Ec; (E) 17 μM FabF with 15 μM ACT$\mathcal{N}$-$\alpha_2$Ec; and (F) 17 μM FabF with 50 μM ACT$\mathcal{N}$-$\alpha_2$Ec. Every 8-th scan is presented for clarity. Data were collected at 280 nm with radial step size 0.002 cm and time delay of 0 s.

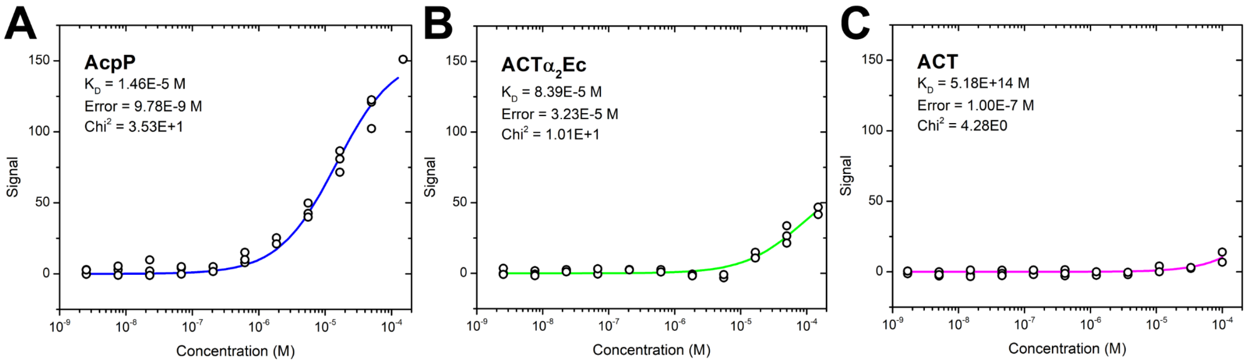


**Figure S17.** SPR affinity fitting curves used in Figure 5D. Affinity fits using all of the collected data (triplicate, except for obvious outliers) for each ACP with FabF KS as a ligand.

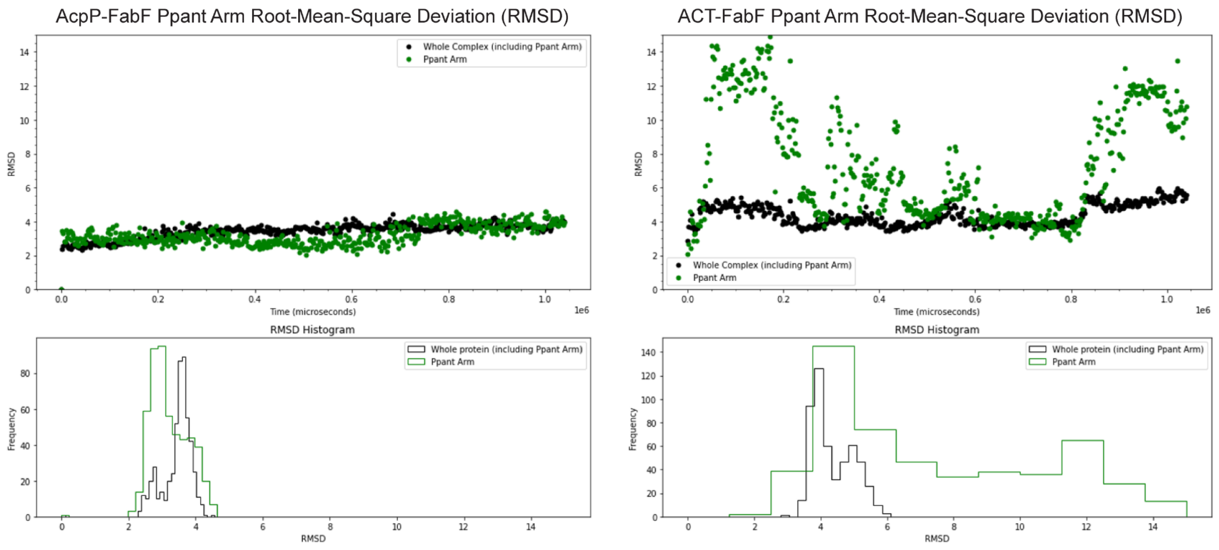


**Figure S18.** Plots of the AcpP-FabF and ACT-FabF complex (black) and Ppant arm (green) root-mean-square deviation over time (top) and histograms of complex and Ppant arm RMSD (bottom).

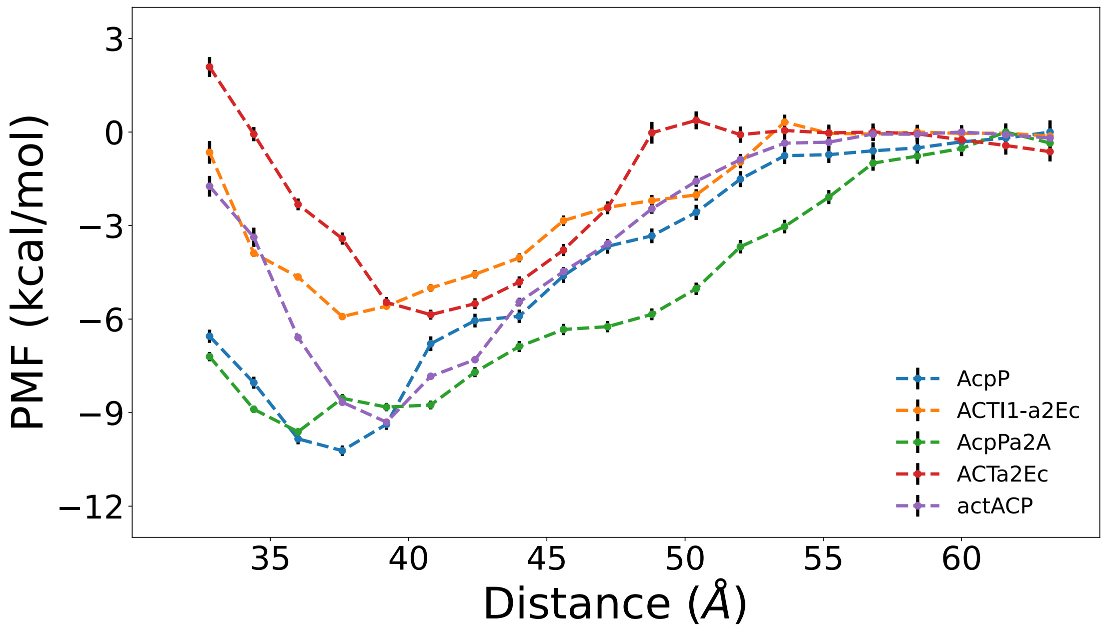


**Figure S19.** Potential of mean force curves for FabF and several ACP structures along a simple center of mass distance coordinate.
